# nanoASM: Long-Read Allele-Specific DNA Methylation Profiling Enables Functional Annotation of Regulatory Noncoding Variants in Human Prostate Tissues

**DOI:** 10.64898/2026.06.17.732357

**Authors:** Yijun Tian, Jodie Wong, Shannon K. McDonnell, Hua Zhong, Lang Wu, Nicholas B. Larson, Brandon J Manley, Liang Wang

## Abstract

Long-read nanopore sequencing enables simultaneous detection of germline variation and native DNA base modifications on individual DNA molecules, providing a unique opportunity to investigate allele-specific epigenetic regulation. Here, we performed whole-genome nanopore sequencing on normal and tumor prostate tissues to characterize differential methylation, methylation entropy, and allele-specific methylation (ASM) associated with noncoding genetic variants. Genome-wide analysis identified extensive cancer-associated differentially methylated regions (DMRs), with hypermethylated DMRs significantly enriched near transcription start sites and transcriptional regulatory regions. Integration with transcriptomic datasets revealed strong inverse relationships between promoter methylation and gene expression, while 5-hydroxymethylcytosine (5hmC) levels positively correlated with transcriptional activity across gene bodies. Using fragment-level methylation patterns enabled by long-read sequencing, we further quantified methylation entropy incorporating both 5mCG and 5hmCG states. Cancer-hypermethylated DMRs exhibited markedly reduced entropy, consistent with clonal fixation of methylation states during tumor progression. Entropy profiling across chromatin annotations demonstrated maximal epigenetic heterogeneity at partially modified enhancer-associated regions. To investigate cis-regulatory genetic effects, we developed a simple ASM framework (nanoASM) that can partition sequencing reads by allelic state and identifies allele-specific DMRs directly from long-read data. Compared with conventional population-level mQTL analysis, ASM demonstrated substantially improved statistical efficiency by leveraging within-individual contrasts and reducing sample-level heterogeneity. Although germline single nucleotide polymorphisms (SNPs) were largely shared between normal and tumor tissues, ASM patterns differed substantially, with tumor-associated ASM regions displaying significantly larger genomic span and stronger allelic methylation differences. Comparative analysis with TCGA prostate mQTL and GTEx prostate eQTL datasets demonstrated substantial concordance between ASM directionality and downstream transcriptional effects, particularly for variants located within DMRs and near transcription start sites. At the IRX4 prostate cancer risk locus, ASM identified an androgen-responsive regulatory domain overlapping AR ChIP-seq and H3K27ac peaks, nominating rs6885084 as a candidate functional variant. At the PSCA locus, ASM anchored by rs4736369 was associated with allele-specific methylation, chromatin activation, transcript abundance, and isoform usage. Together, these findings establish nanopore-based ASM analysis as a powerful approach for resolving functional noncoding variants and their regulatory domains they control in prostate cancer.

## Introduction

The completion of the Human Genome Project (HGP) marked a major milestone in biomedical research, providing a nearly complete reference sequence of the human genome.^1-3^ Although the vast majority of the genome is identical among individuals, phenotypic diversity is largely driven by single nucleotide polymorphisms (SNPs).^4,5^ Deciphering the functional impact of these genetic variants, however, remains a formidable challenge.^6-8^ Genome-wide association studies (GWAS) have identified thousands of SNPs associated with complex traits and diseases, yet most reside in noncoding regions where their biological interpretation remains elusive.^9,10^ To link genetic associations with molecular function, expression quantitative trait locus (eQTL) mapping has been widely applied to connect germline variants with gene expression,^11,12^ revealing that SNPs can modulate transcription by altering enhancer activity,^13^ promoter accessibility,^14^ or chromatin organization.^15^ Nevertheless, transcript-level associations alone provide only a partial view of regulatory complexity, as epigenetic modifications — particularly DNA methylation — play a pivotal role in shaping transcriptional outcomes.

Methylation quantitative trait locus (mQTL) analysis extends these efforts by identifying variants that influence DNA methylation patterns, revealing how sequence differences can alter local chromatin states and gene regulation.^16-18^ Despite its strong functional implications, current mQTL approaches are constrained by several methodological limitations. Although mQTL studies have identified numerous statistically significant associations, the majority rely on microarray-based methylation data in which the signal corresponds to isolated single-CpG sites rather than broader functional regions.^19-21^ Combined with the effects of linkage disequilibrium (LD), this produces a large number of associations distributed broadly across the genome, many of which offer limited power to prioritize functionally relevant variants. ^18^

One strategy to more effectively pinpoint functional noncoding variants is to evaluate their regulatory potential by examining the local methylation pattern flanking each allele.^22^ Such allele-specific information can indicate whether a variant perturbs proximal epigenetic states and thereby influences regulatory activity — but this requires the simultaneous quantification of base modifications and variant alleles within the same DNA molecules. Bisulfite-based methods combined with microarray or next-generation sequencing (NGS) technologies have enabled genome-wide methylation profiling, yet they carry inherent limitations. Bisulfite conversion destroys allelic information at cytosine-containing SNPs, complicating variant detection. Short-read sequencing further restricts the analysis of co-methylation patterns among distal CpG sites, and PCR amplification can distort native methylation ratios. Critically, bisulfite-based approaches cannot distinguish 5-methylcytosine (5mC) from its oxidized derivative 5-hydroxymethylcytosine (5hmC)^23^, which serves as an intermediate in active demethylation^24,25^ and plays independent regulatory roles in transcription and development.^26^

To overcome these limitations, we employed nanopore long-read sequencing, which directly detects base modifications and sequence variation on the same DNA molecules without bisulfite conversion. We aggregated read-level methylation by the allele carried on each read, obtaining one methylation estimate per allele per individual (reference vs. alternative), and compared allele-specific methylation (ASM) differences across individuals. Conceptually, this framework operates at the allele level rather than on bulk sample averages: heterozygous individuals contribute two allele-specific observations under a shared genetic background, while homozygous individuals contribute genotype-dosage information. This allele-resolved design reduces the influence of sample-level heterogeneity relative to conventional population-level mQTL analysis, thereby increasing effective power to detect *cis* effects — particularly for common variants.

## Methods

### Cohort information and case selection

As demonstrated in our previous study, a germline risk variant can induce allele-specific epigenetic differences that are substantially altered during tumorigenesis.^14^ This observation led us to hypothesize that inherited genetic variation may contribute to cancer-associated epigenetic remodeling and that such effects can be systematically identified through allele-specific methylation analysis. Therefore, in addition to normal prostate tissues, we included prostate cancer samples to investigate how genetic regulation of DNA methylation is maintained, altered, or amplified in the cancer state. Prostate genomic DNA and tissue samples used for nanopore sequencing were collected from two institutions: Mayo Clinic (dbGaP accession phs000985.v2.p1, "Functional Significance of Prostate Cancer Risk-SNPs") and Moffitt Cancer Center ("Total Cancer Care Biobanking"). The informed-consent process was completed with all participants, and the study was approved by the Institutional Review Boards of both institutions.

For the Mayo Clinic cohort, archived genomic DNA (N = 30) was used directly for nanopore sequencing library preparation. Prostate tissue was retrieved from archived fresh-frozen collections obtained from patients who underwent radical prostatectomy or cystoprostatectomy. For each normal prostate tissue sample, H&E-stained slides were reviewed and required to meet the following criteria: (1) absence of cancer cells; (2) absence of benign prostatic hyperplasia; (3) high representation of normal prostatic epithelial glands (≥40% of all cells); and (4) low fraction of lymphocytic infiltrate (≤20% of all cells).

For the Moffitt Cancer Center cohort, primary prostate tumor (N = 15) and paired adjacent normal tissue (N = 15) were obtained from patients with prostate cancer receiving surgical treatment under protocols approved by the Moffitt Cancer Center Institutional Review Board (Total Cancer Care protocol [TCC], MCC# 50468; Advarra IRB Pro00000971).

To ensure a uniform genetic background, all patients included in this study are self-reported as Caucasian male.

### Tissue processing and genomic DNA extraction

For the Mayo Clinic cohort, tissue preparation and genomic DNA extraction procedures have been described previously ^27^. For the Moffitt Cancer Center cohort, fresh-frozen tissue samples were pulverized, and genomic DNA was extracted using the QIAamp DNA Blood Kit (Qiagen). Prior to proteinase K digestion, RNase A was added to the tissue lysate to ensure thorough RNA removal. DNA quality was assessed by the 260/280 absorbance ratio and total DNA yield.

### Nanopore library preparation and multiplexed sequencing

Prior to library preparation, genomic DNA samples were sheared to 8–10 kb fragments using a g-TUBE (Covaris). Sheared DNA was end-repaired and used for library preparation with the Native Barcoding Kit (SQK-LSK114). The multiplexed library was loaded onto an R10.4.1 PromethION flow cell (FLO-PRO114M) for long-read sequencing data acquisition. After sequencing, raw signal was base-called using Dorado with the model dna_r10.4.1_e8.2_400bps_sup@v5.0.0 to infer both genomic sequence and CpG base modification probabilities. Base calling produced a BAM file per sample containing aligned reads with per-CpG modification probability tags, which were used for all downstream analyses.

### Nanopore sequencing quality control

Quality control was applied at two levels. First, read-level quality was assessed by the median per-read quality score, and samples with a median score below Q20 were excluded. Second, tissue composition was evaluated by CpG methylation deconvolution.^28^ To ensure that each sample was dominated by prostate epithelial cells, we calculated a purity ratio defined as the fraction of prostate epithelium divided by the fraction of the second most abundant cell type. Samples were retained only if they met both criteria: prostate epithelium fraction >40% and purity ratio ≥1.5. Applying these filters, three samples from the Moffitt cohort (Cancer.P10, Normal.P06, Normal.P08) and eight samples from the Mayo Clinic cohort (s86, s460, s307, s160, s315, s424, s1139, s112) were excluded. A summary of sequencing metrics for all retained samples is provided in Supplementary Table 1.

### Copy number variation analysis

Copy number variation (CNV) analysis was performed using CNVkit^29^ to assess genome-wide DNA integrity. Because this analysis relies solely on alignment depth, all samples passing per-read quality thresholds were included regardless of methylation deconvolution status. A pooled reference was constructed by combining BAM files from all normal samples across both cohorts to derive per-bin copy number estimates. This reference was then applied to all prostate tumor samples. CNVkit was run in whole-genome sequencing mode (--method wgs), with very low-coverage bins excluded prior to segmentation (--drop-low-coverage). Genome-wide CNV patterns were visualized as a sample-wise heatmap, and scatter plots were used to illustrate focal copy number gains in selected samples.

### Differential methylation region (DMR) identification between tumor and normal tissue

Following Dorado base calling, CpG-level methylation profiles were summarized into BEDGRAPH format using modkit (https://github.com/nanoporetech/modkit). CpG sites were retained if covered by ≥10 reads, and the resulting profiles were used as input for de novo DMR detection with metilene.^30^ Significant DMRs were defined by a Bonferroni-adjusted q-value ≤0.05 and an absolute tumor–normal methylation difference ≥10%. For downstream Genomic Regions Enrichment of Annotations Tool (GREAT) analysis, significant DMRs were stratified into hypermethylated and hypomethylated subsets. All tested genomic regions, irrespective of statistical significance, were used as the background reference for GREAT analysis.

### Methylation entropy analysis

To evaluate epigenetic heterogeneity in prostate tissue, we performed methylation entropy analysis. Methylation entropy quantifies the diversity of methylation patterns by enumerating all possible methylation states across a fixed number of adjacent CpG sites. Entropy profiles were calculated for every set of three consecutive CpG sites using the modkit entropy function. An entropy value of zero indicates a uniform methylation pattern across reads, whereas higher values reflect increased heterogeneity and more complex methylation configurations. Results were stratified by the genomic context of the central CpG site (within or outside CpG islands) and by the genomic distance spanned by each CpG triplet. Given the large proportion of zero-entropy observations, downstream analyses were restricted to CpG triplets with non-zero entropy values.

### Variant calling and read-level allele-specific methylation analysis

Variant calling was performed per sample using PEPPER-Margin-DeepVariant, a pipeline^31^ designed to improve SNP calling accuracy in homopolymer regions. Called variants were annotated against dbSNP build 153, and annotated SNPs were filtered by variant quality score (QUAL ≥10) and variant allele fraction (VAF ≥0.1). Filtered VCF files were merged across samples and further refined by minimum allele count (--min-ac 3), heterozygous sample allele depth (≥3 heterozygous samples with FORMAT/DP ≥10, FORMAT/AD ≥5), and post-merge variant quality (QUAL ≥30). Only biallelic SNPs were retained for allele-specific methylation (ASM) analysis.

To prepare methylation signals for allele-level comparisons, individual BAM files were processed with the modkit call-mods subcommand, which binarizes per-CpG modification probabilities to 0% or 100% using sample-specific thresholds. This step ensures consistent methylation calls across samples. Processed BAM files were then merged with sample identifiers encoded in the read group (RG) tag.

Each input variant’s coordinate and allele information were used to label reads in the merged BAM with a variant allele tag (VA), indicating whether each read carries the reference or alternative allele. Labeled reads were processed with modkit pileup with sample (RG) and allele (VA) partitioning enabled, generating per-allele BEDGRAPH files. DMRs between reference and alternative allele groups were identified de novo using metilene with default settings. Prior to segmentation, samples with ≥90% missing CpG methylation values across a given region were excluded to prevent instability from sparse coverage. The full script developed has been deposited in the nanoASM GitHub repository (https://github.com/Yijun-Tian/nanoASM).

## Results

### Quality Control and Initial Characterization of Nanopore Long-Read Sequencing Data

Following nanopore sequencing, we performed quality control at multiple levels. Per-read quality score distributions across all samples from both the Mayo Clinic and Moffitt cohorts showed peaks well above Q20, consistent with the expected performance of the R10.4.1 PromethION platform (**Figure S1A**). Notably, within the Moffitt cohort, prostate cancer tissue samples exhibited significantly lower per-read quality scores compared with their paired normal counterparts (KS-test p < 2.2×10^-16^; **Figure S1B**). We hypothesize that this reflects a greater diversity of complex or atypical base modifications in cancer cells, which may reduce base calling confidence. Consistent with this interpretation, our previous work^22^ demonstrated that the concordance between nanopore and bisulfite sequencing is systematically lower in cancer cell lines than in normal cell lines, suggesting a similar underlying phenomenon.

To characterize cellular heterogeneity in the sequencing data, we performed cell-type deconvolution using nanopore-derived CpG methylation profiles referenced against the ATLAS reference pane.^28^ The inferred cell-type composition confirmed that most samples were dominated by prostate epithelial cells (Figure S1C). However, three samples from the Moffitt cohort (Cancer.P10, Normal.P06, Normal.P08) and eight samples from the Mayo Clinic cohort (s86, s460, s307, s160, s315, s424, s1139, s112) - highlighted in red in Figure S1C — showed insufficient prostate epithelial fractions and were excluded from downstream analyses.

Copy number variation (CNV) analysis was performed on both normal and tumor prostate tissues using alignment depth information. Despite most tumors being derived from early-stage surgical resections, cancer tissues exhibited substantially greater copy number variation than matched normal tissues (**Figure S2A**). For instance, a log_2_ ratio scatter plot of chromosome 8 in patient 06 revealed the characteristic pattern of 8p loss and 8q gain/amplification, a hallmark of prostate cancer (**Figure S2B**).

### Genome-wide Identification of Differentially Methylated Regions in Prostate Cancer

Using the high-quality nanopore long-read sequencing data, we performed de novo identification of differentially methylated regions (DMRs) between normal and cancer tissues. After annotating these DMRs using GREAT analysis, we observed that DMRs exhibiting higher methylation in cancer intersected significantly more transcription start site (TSS)–proximal regions compared with DMRs that were more highly methylated in normal tissues (Figure 1A).

When further annotating DMRs to known human genes, we found that 228 cancer-hypermethylated DMRs and only three normal-hypermethylated DMRs did not overlap with any gene regions (**Figure 1B**). Functional annotations using Gene Ontology (GO) Molecular Function revealed that cancer-hypermethylated DMRs were significantly enriched in genomic regions associated with transcriptional regulation, including sequence-specific DNA binding and RNA polymerase II regulatory regions. In contrast, DMRs hypermethylated in normal tissues were enriched in regions associated with ion channel receptor activity, aminotransferase activity, and related molecular functions (**Figure 1C**). Based on the DMR-to-gene mapping, we then leveraged RNA expression profiles from the TCGA prostate cancer cohort (PRAD) to examine the relationship between gene expression and DMR methylation differences. In both the PRAD normal and cancer subgroups, the strongest correlations between gene expression and DNA methylation were consistently enriched near transcription start sites, particularly for DMRs that were hypermethylated in cancer tissues. This observation is consistent with the transcription regulatory functions identified in our functional annotation analysis (**Figure 1D**). To highlight representative DMR–gene associations, we visualized the aggregated methylation levels of the most significant DMRs (p ≤ 0.00001) in the Moffitt cohort (**Figure 1E**), alongside the corresponding gene expression levels in matched tumor-normal pairs from the TCGA prostate cancer cohort (**Figure 1F**). Several genes previously reported to exhibit recurrent methylation alterations in prostate cancer were highlighted in the heatmap, including RASSF1,^32,33^ TWIST1,^34,35^ AOX1,^36,37^ GPX3,^38-40^ DLEC1,^41^ ZNF154,^42^ and WNT5A.^43,44^ As expected, the genes showed over-expression in TCGA-PRAD tumor tissue mostly showed hypomethylation in Moffitt tumor tissues.

**Figure 1.**
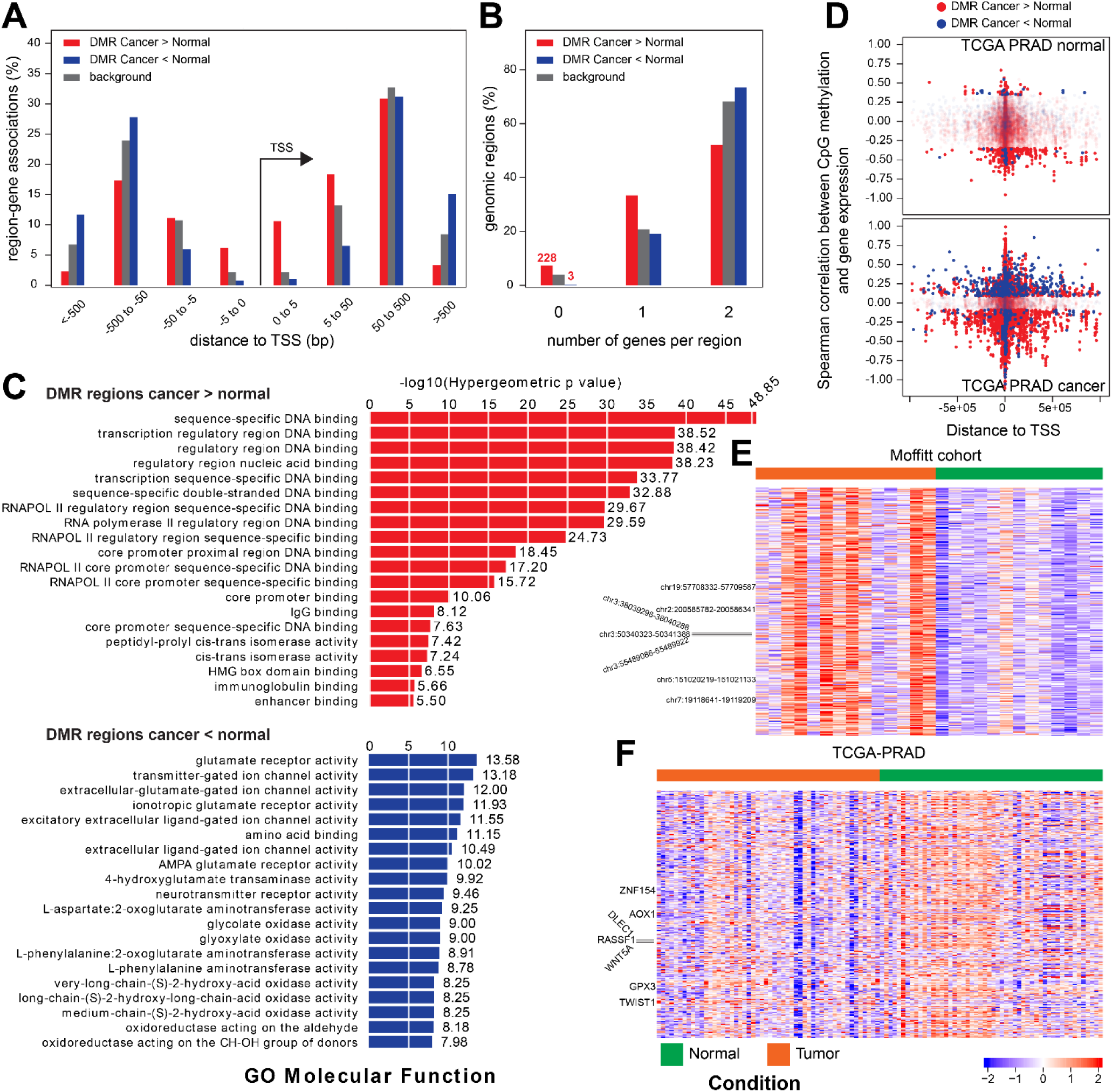
Characterization of differentially methylated regions between prostate cancer and normal tissue. (**A**) Distribution of region–gene associations by distance to the nearest transcription start site (TSS) for hypermethylated DMRs (Cancer > Normal, red), hypomethylated DMRs (Cancer < Normal, blue), and background genomic regions (gray). (**B**) Distribution of the number of genes associated per DMR for hypermethylated (red), hypomethylated (blue), and background (gray) regions. (**C**) Gene Ontology (GO) Molecular Function enrichment analysis of hypermethylated (top, red) and hypomethylated (bottom, blue) DMRs, performed using GREAT. Bar lengths represent −log_10_(hypergeometric p-value); values are labeled at the right of each bar. (**D**) Scatter plots of Spearman correlation between CpG methylation and gene expression as a function of distance to TSS, derived from TCGA PRAD normal (top) and cancer (bottom) samples. Red dots represent hypermethylated DMRs (Cancer > Normal) and blue dots represent hypomethylated DMRs (Cancer < Normal). (**E**) Heatmap of DMR methylation levels across Moffitt cohort samples (normal, green; tumor, orange). Selected DMR coordinates are labeled on the left. DMRs clearly stratify tumor from normal samples, with hypermethylated regions (red) predominantly enriched in cancer and hypomethylated regions (blue) enriched in normal tissue. (**F**) Heatmap of expression levels of DMR annotated genes in TCGA PRAD samples (normal, green; tumor, orange). Selected known prostate cancer–associated genes (ZNF154, AOX1, DLEC1, RASSF1, WNT3A, GPX3, TWIST1) are labeled on the left. Color scale represents methylation z-score (−2 to +2).

### RNA expressions associated with epigenetic profiling detected in nanopore DNA sequencing

Current models suggest that DNA modifications act as stable and functional epigenetic markers that are strongly correlated with gene expression. However, most approaches used to investigate DNA modifications rely on capture-based methods, and many studies have been conducted primarily in mouse models or in vitro systems.^45-47^ By enabling single-base resolution detection of both 5mCG and 5hmCG in native genomic DNA, nanopore sequencing in our cohort provides an opportunity to re-evaluate these established relationships in human prostate tissues.

We first visualized the average 5mCG (Figure 2A) and 5hmCG (Figure 2B) modification levels across genes grouped by their expression levels based on GTEx prostate tissue data (n = 231). Notably, the average 5mCG level in transcription start site (TSS) regions decreased with increasing gene expression (Figure 2C), whereas the 5hmCG level in gene body regions increased with gene expression (Figure 2D). These patterns suggest that the two epigenetic modifications may play distinct regulatory roles across different genomic regions.

**Figure 2.**
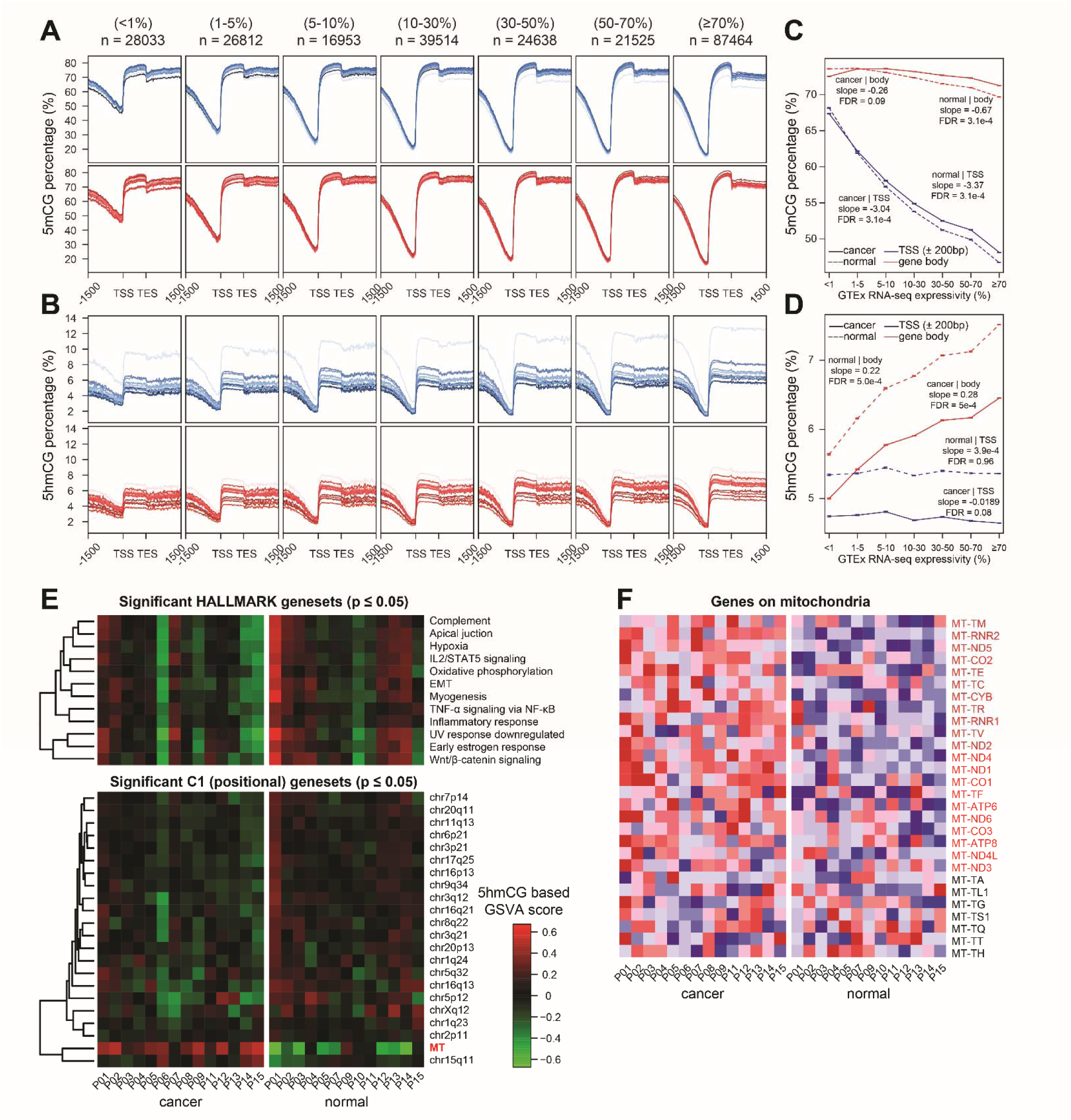
Genome-wide 5mCG and 5hmCG profiles across gene bodies stratified by expression level. **(A)** Meta-gene profiles of 5mCG percentage across gene bodies (TSS to TES, ±1500 bp flanking) for normal (blue, top) and cancer (red, bottom) prostate tissue samples, stratified by GTEx RNA-seq expression expressivity bins (<1%, 1–5%, 5–10%, 10–30%, 30–50%, 50–70%, ≥70%). The number of genes in each bin is indicated above each panel. **(B)** Meta-gene profiles of 5hmCG percentage across gene bodies for normal (blue, top) and cancer (red, bottom) prostate tissue samples, using the same expression percentile stratification as in (A). **(C)** Spearman correlation between 5mCG levels and gene expression percentile at the TSS (±200 bp, solid lines) and across the gene body (dashed lines) for cancer (red) and normal (blue) tissue. **(D)** Spearman correlation between 5hmCG levels and gene expression percentile at the TSS (±200 bp, solid lines) and across the gene body (dashed lines) for cancer (red) and normal (blue) tissue. **(E)** Heatmap of 5hmCG-based GSVA enrichment scores for significant HALLMARK gene sets (top, p ≤ 0.05) and C1 positional gene sets (bottom, p ≤ 0.05) across Moffitt cancer and normal samples. Red indicates positive enrichment and green indicates negative enrichment. **(F)** Heatmap of 5hmCG levels across individual mitochondrial genes in cancer (left) and normal (right) Moffitt samples. Mitochondrial genes in the leading edge are labeled in red while those are not labeled in black.

Given that 5hmCG has been reported to be strongly associated with transcriptional activity, we further performed gene set enrichment analysis (GSEA) using gene-level aggregated 5hmCG measurements across the HALLMARK (h) and positional (c1) gene set collections. Interestingly, multiple hallmark pathways and autosomal positional gene sets exhibited higher 5hmCG levels in normal tissues compared with cancer tissues (Figure 2E). In contrast, mitochondrial gene sets displayed significantly higher 5hmCG levels in cancer tissues relative to normal tissues (Figure 2F).

### Entropy-based characterization of CpG methylation landscapes

In addition to analyzing the 5hmCG and 5mCG information on a per CpG-site basis, we adopted methylation entropy in evaluating the methylation diversity from the nanopore sequencing data. Methylation entropy recognizes epigenetic diversities defined by combinatory modifications including 5hmCG, 5mCG and CG without modifications to describe the epigenetic diversity in each set of fixed CpG site triplets (**Figure 3A**). At the genome-wide level, regional entropy was comparable between normal and cancer samples, indicating a highly similar background distribution of CpG methylation configurations (**Figure 3B**). Consistent with this, no significant differences were observed after stratifying by CpG island status (**Figure S3A**) or gene-centric annotations (**Figure S3B**). However, when focusing on normal–cancer DMRs, a directional difference was observed: regions that become hypermethylated in cancer (DMR normal < cancer) showed lower entropy in cancer, whereas hypomethylated regions (DMR cancer < normal) did not show a consistent change (**Figure 3B**).

**Figure 3.**
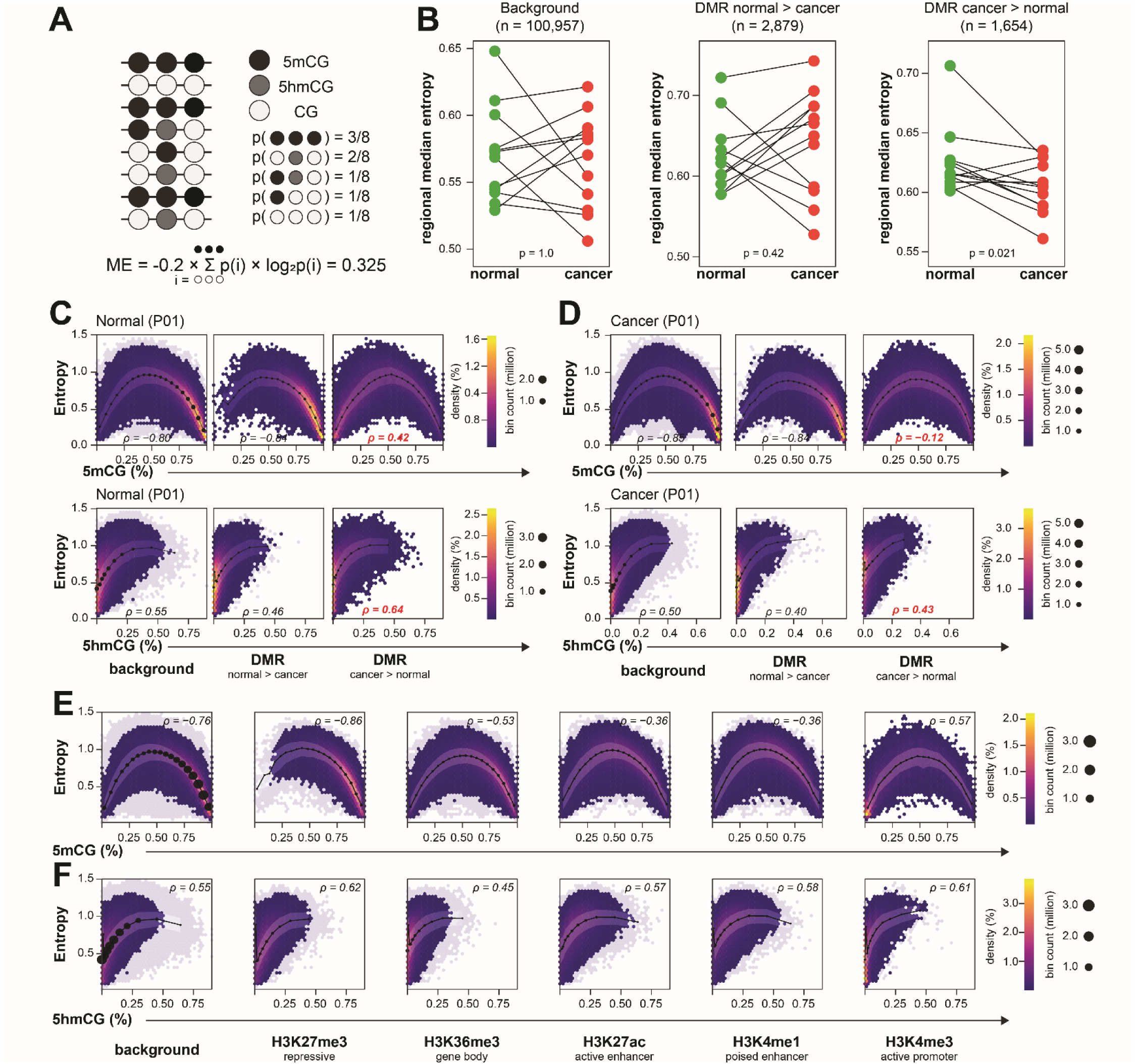
Methylation entropy characterizes epigenetic heterogeneity across DMRs and chromatin states. (**A**) Schematic illustrating the calculation of methylation entropy (ME) for a set of three adjacent CpG sites. Each row represents a distinct methylation pattern observed across sequencing reads, with filled circles (black: 5mCG, gray: 5hmCG, white: unmodified CG) indicating the modification state at each position. The probability of each pattern is enumerated, and ME is computed as −0.2 × Σ p(i) × log₂p(i). (**B**) Paired dot plots comparing regional median methylation entropy between matched normal (green) and cancer (red) samples from the Moffitt cohort, stratified by genomic context: background CpG triplets (n = 100,957; left), hypomethylated DMRs in cancer (DMR normal > cancer, n = 2,879; middle), and hypermethylated DMRs in cancer (DMR cancer > normal, n = 1,654; right). (**C**) Hexbin density scatter plots of methylation entropy versus 5mCG percentage (top row) and 5hmCG percentage (bottom row) for the normal tissue sample from patient P01, stratified by genomic context: background (left), DMR normal > cancer (middle), and DMR cancer > normal (right). Color scale indicates bin count density. Spearman correlation coefficients (ρ) are shown for each panel; values highlighted in red indicate significant differences from the background correlation. (**D**) Equivalent hexbin density scatter plots as in (C) for the matched cancer tissue sample from patient P01. (**E**) Hexbin density scatter plots of methylation entropy versus 5mCG percentage stratified by histone modification chromatin state, including background, H3K27me3 (repressive), H3K36me3 (gene body), H3K27ac (active enhancer), H3K4me1 (poised enhancer), and H3K4me3 (active promoter). Spearman ρ values are shown for each context. (**F**) Hexbin density scatter plots of methylation entropy versus 5hmCG percentage across the same chromatin state categories as in (**E**).

To further characterize this pattern, we examined the relationship between entropy and modification levels in a representative sample (**Figure 3C–D**). In regions corresponding to DMR normal < cancer, the correlation between 5mCG and entropy was positive (ρ = 0.42) in normal tissue but became negative (ρ = -0.12) in cancer. This shift indicates a change in how CpG methylation levels relate to local configuration variability. In the same regions, entropy showed a consistent positive association with 5hmC, suggesting that variation in entropy tracks more closely with 5hmCG than with 5mCG. In contrast, background region sets showed more stable or weaker relationships. Consistent with their genomic distribution (**Figure 1C**), hypermethylated DMRs are enriched near transcription start sites and regulatory elements, whereas hypomethylated DMRs are more broadly distributed. Together, these results describe a context-dependent change in the relationship between methylation level and entropy in cancer.

Finally, we assessed whether CpG methylation and entropy profiles alone could recapitulate underlying chromatin context (**Figure 3E–F**). Using nanopore-derived signals, regions stratified by histone modification annotations from normal prostate tissue displayed distinct methylation–entropy patterns consistent with their regulatory states. More specifically, repressive (H3K27me3) and gene body (H3K36me3) chromatin states displayed increased density associated with strong 5mCG signals (>0.75), weak 5hmCG signals (<0.1), and low methylation entropy (0-0.5). Promoter regions marked by H3K4me3 showed low levels of both 5mCG (<0.25) and 5hmCG (<0.1) and correspondingly low entropy (0-0.1). Interestingly, enhancer-associated histone modifications (H3K27ac and H3K4me1) exhibited intermediate 5mCG (0.25-0.75) and high 5hmCG (0.1-0.25) levels together with high entropy (0.5-1.0). Across all contexts, entropy consistently peaked at intermediate modification levels on 5mCG, with reduced entropy observed toward fully methylated or unmethylated states (**Figure 3C–F**). This intermediate-state dependence was reproducible across samples and genomic annotations, indicating that CpG configuration diversity is maximized in partially modified regions.

Consistent patterns were also observed in a cancer type–specific chromatin accessibility framework (Figure S3C), where regions defined by accessibility across tumor types exhibited the same enrichment of high entropy at intermediate methylation levels. Together, these results highlight the intermediate methylation state as a key determinant of local CpG configuration variability and support the use of methylation–entropy profiles as informative proxies for underlying chromatin state.

### Theoretical Evaluation of Statistical Efficiency Between ASM and Population-Based mQTL testing

To improve sensitivity for detecting cis-regulatory methylation effects, we implemented an allele-specific methylation (ASM) framework based on within-individual allelic contrasts. In heterozygous individuals, both alleles are measured within the same cellular and experimental context, allowing shared sample-level variation to be removed through paired comparison. As a result, the residual variation is primarily driven by technical noise, leading to a substantial reduction in variance relative to population-level analyses.

This property contrasts with traditional mQTL approaches, which rely on between-individual comparisons of bulk methylation levels and are therefore more susceptible to sample-level heterogeneity and measurement variability. In addition, ASM analysis leverages sequencing reads that capture both allele identity and CpG methylation states, often aggregating information across multiple CpGs within a local genomic region. This aggregation further stabilizes methylation estimates and reduces effective measurement noise. The advantage is particularly pronounced in long-read sequencing data, where haplotype information and contiguous CpG methylation patterns are directly observed within individual reads.

The efficiency gain of ASM increases with the proportion of heterozygous individuals in the cohort, as more samples contribute informative allelic contrasts. Under a Gaussian approximation, and defining bulk methylation as the average of allele-specific measurements, the relative statistical efficiency of ASM compared to population-level testing can be approximated as

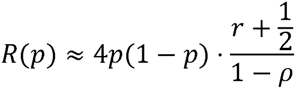

where p is the allele frequency, r is the ratio of sample-level to technical variance, and ρ represents the correlation of allele-specific technical errors within individuals. This formulation yields more conservative but more realistic conditions under which ASM provides improved statistical power.

In particular, for common variants (p≈0.5) and settings with substantial sample-level variability (large r), ASM can achieve markedly higher efficiency than population-based mQTL analysis (Figure S4A). Consequently, comparable statistical power can be obtained with substantially smaller sample sizes (Figure S4B–C). Full derivations and simulation results are provided in Supplementary Note 1.

### Allele-specific Methylation Identification from Nanopore Long-read DNA sequencing

Given the evidence of the improved statistical power in the ASM framework over traditional mQTL approaches, we sought to implement a pipeline to identify the read-level allele-specific methylation region from Nanopore sequencing BAM files with base modification tags. For each heterozygote variant, we labeled and grouped the nanopore long reads according to the allele they carried. After labeling, we performed the de-novo DMR identification within the reference and alternative allele group (**Figure 4A**). Given any variant, we were able to use this simple method to identify differential methylation domains among the read spanning regions across the whole genome. From 4,076,763 and 5,788,657 input variants, we identified 406,011 (**Figure 4B**) and 391,918 (**Figure 4C**) allele-specific methylation events within the tumor and normal subgroups respectively. From these findings, we further defined shared ASM events as those associated with the same SNP and overlapping differential methylation regions between tumor and normal. To evaluate the correlation in effect size between the ASM and mQTL method, we intersected the ASM findings with the Pancan-meQTL results from TCGA prostate cancer cohort by the SNP id and the DMR-to-probe inclusion. We found majority of overlapped associations showed the same trend of allelic effect (**Figure 4D-4E**). Interestingly, although we observed large overlaps in the input SNPs between normal and tumor subgroups (**Figure 4F**), SNPs identified with ASM (**Figure 4G**) and the relevant ASM (**Figure 4H**) overlapping is relatively small. We further compared the genomic size and the absolute methylation difference of the DMR between normal and cancer subgroups and found that cancer ASM showed significantly larger DMR size (**Figure 4I**) and stronger allelic methylation difference (**Figure 4J**) than normal ASM. We had also explored whether allele-related CpG site change could influence DMR methylation direction. We observed that when a SNP is in proximity to a DMR, allele forming a CpG site is associated with hypermethylation, especially in those ASM events identified in normal tissue (**Figure 4K**).

**Figure 4.**
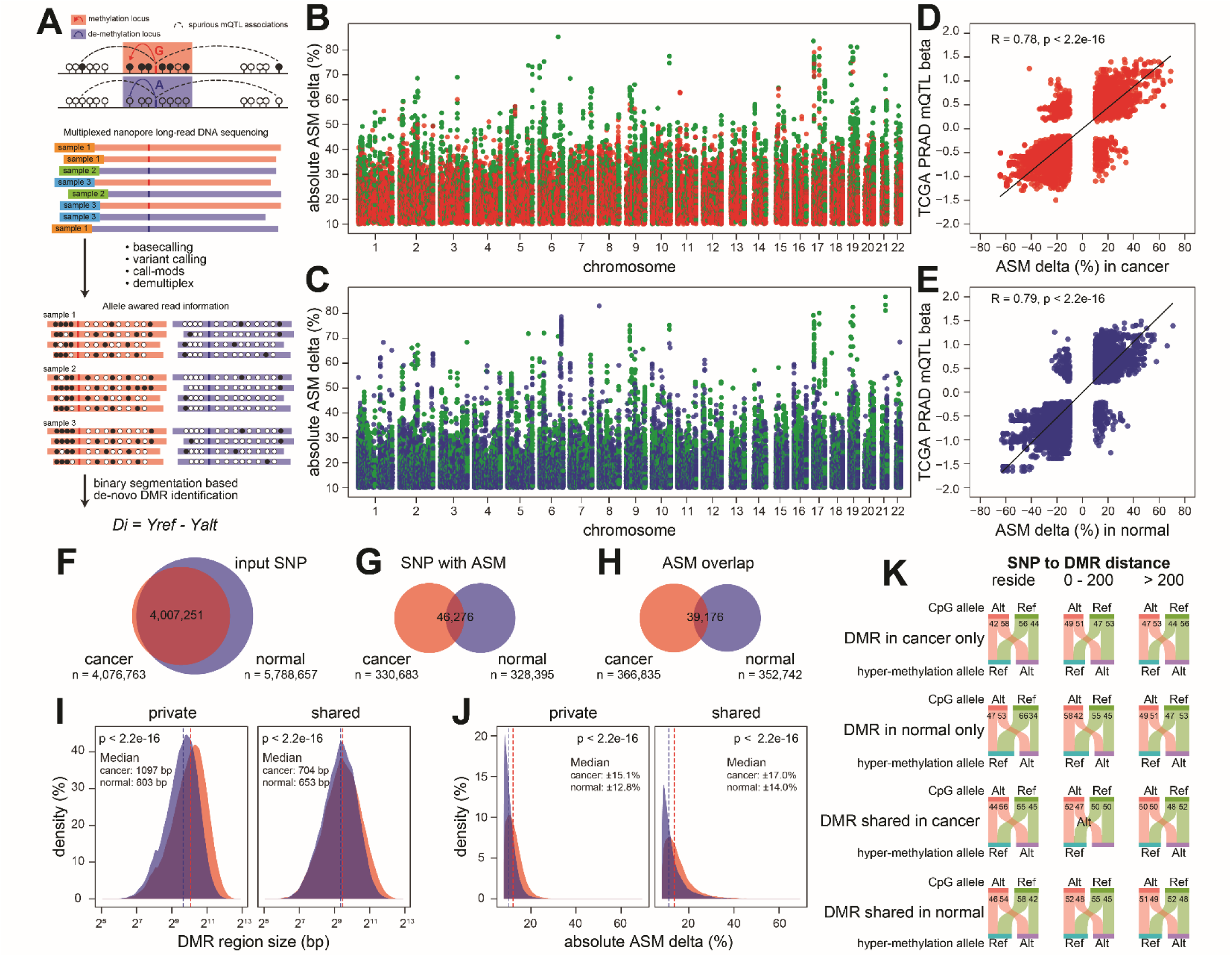
Genome-wide allele-specific methylation analysis using nanopore long-read sequencing. **(A**) Schematic of the allele-specific methylation (ASM) analysis framework. Multiplexed nanopore long-read sequencing simultaneously captures base modifications and sequence variants on individual DNA molecules. After basecalling, variant calling, call-mods processing, and demultiplexing, reads are labeled by the allele they carry (reference or alternative). Per-allele methylation signals are aggregated across samples, and de novo DMRs between reference and alternative allele groups are identified by binary segmentation. The ASM delta (Di = Yref − Yalt) quantifies the methylation difference between alleles. (**B-C**) Genome-wide Manhattan-style scatter plot of absolute ASM delta (%) for all significant ASM-DMRs identified in prostate cancer (**B**) and normal (**C**) tissue, plotted by chromosomal position. Green dots represent ASM-DMRs unique to cancer; red dots represent ASM-DMRs shared between cancer and normal tissue; Blue dots represent ASM-DMRs unique to normal tissue. (**D**) Scatter plot comparing ASM delta (%) in cancer tissue against TCGA PRAD mQTL beta coefficients. (**E**) Scatter plot comparing ASM delta (%) in normal tissue against TCGA PRAD mQTL beta coefficients. (**F**) Proportional Venn diagram showing the overlap of input SNPs (n = 4,007,251) between cancer (red, n = 4,076,763) and normal (blue, n = 5,788,657) tissue analyses. (**G**) Proportional Venn diagram showing the overlap of SNPs (n = 46,276) with significant ASM between cancer (red, n = 330,683) and normal (blue, n = 328,395) tissue. (**H**) Proportional Venn diagram showing the overlap of ASM-DMR genomic intervals (n = 39,176) between cancer (red, n = 366,835) and normal (blue, n = 352,742) tissue. (**I**) Density distributions of ASM-DMR region size (bp, log₂ scale) for private (left) and shared (right) ASM-DMRs in cancer (red) and normal (blue) tissue. Dashed vertical lines indicate median values. (**J**) Density distributions of absolute ASM delta (%) for private (left) and shared (right) ASM-DMRs in cancer (red) and normal (blue) tissue. (**K**) Stacked Sankey plots showing the allelic distribution in CpG site (Alt allele, teal; Ref allele, pink) and the direction of allelic-methylation (hyper-methylation allele: Ref or Alt) for ASM-DMRs stratified by tissue context (cancer only, normal only, shared in cancer and shared in normal) and by SNP-to-DMR distance (reside: SNP within DMR; 0–200 bp: proximal; >200 bp: distal). Numbers within bars indicate the percentage of ASM-DMRs in each category. SNPs residing within or proximal to DMRs show a strong tendency for the alternative allele to be the hypermethylated allele, while this asymmetry diminishes with increasing SNP-to-DMR distance.

### Consistency of ASM across mQTL and eQTL signals

Having observed the high consistency between ASM and mQTL signals (**Figure 4D–E**), we next sought to better understand where and why these signals differ, and to extend this comparison to eQTL effects. To this end, we first defined discrepancy between mQTL and ASM based on the agreement between DMR-level methylation differences (DMR delta) and mQTL effect sizes (**Figure 5A**). For each SNP–DMR (ASM) and SNP-CpG (mQTL) pair, discrepancy was defined as the inconsistency in direction between allele-specific regional methylation delta and probe-level mQTL effect size. Interestingly, discrepancy between TCGA PRAD mQTLs and cancer-derived ASM showed a clear distance-dependent pattern (**Figure 5B**). Discrepancy decreased sharply from very short SNP–CpG distances (<50 bp) to intermediate distances (∼75–150 bp), and then gradually increased with increasing distance, reaching higher levels in distal regions (>2000 bp). In contrast, discrepancy decreased monotonically with increasing CpG density, with higher discrepancy observed in CpG-sparse regions and lower discrepancy in CpG-dense regions (density > 0.07-0.08).

**Figure 5.**
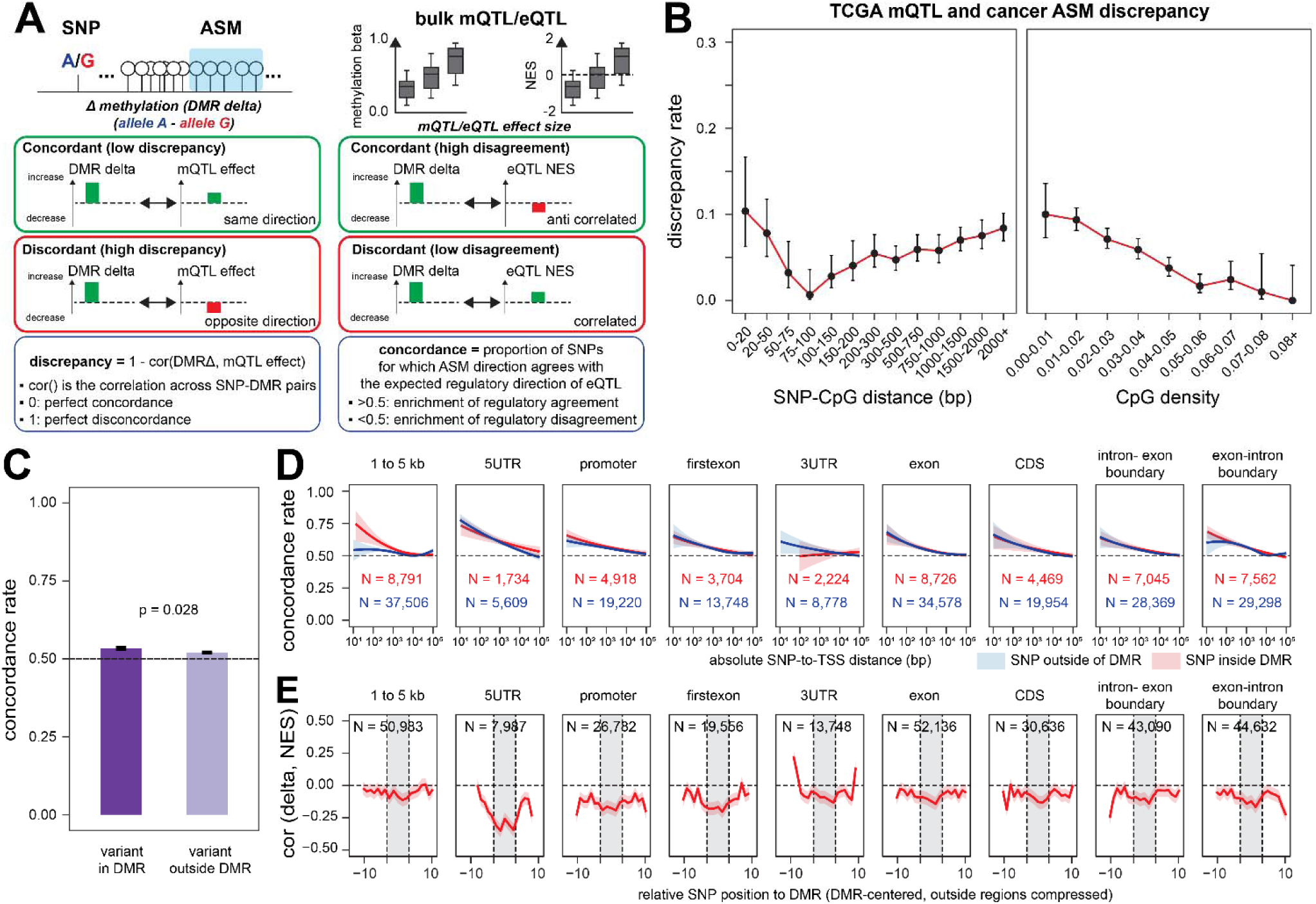
Concordance and discrepancy between nanopore ASM and bulk mQTL/eQTL effect sizes. **(A)** Schematic defining the metrics used to compare ASM-DMR delta values with bulk mQTL and eQTL effect sizes. The ASM delta (Δ methylation = allele A − allele G) is compared with the mQTL methylation beta coefficient or eQTL normalized enrichment score (NES). **(B)** TCGA mQTL and cancer ASM discrepancy rate as a function of SNP-to-CpG distance (left, binned in bp) and local CpG density (right). **(C)** Bar plot comparing the eQTL concordance rate between SNPs residing within ASM-DMRs (dark purple, N = variant in DMR) and SNPs outside ASM-DMRs (light purple, N = variant outside DMR). **(D)** Concordance rate as a function of absolute SNP-to-TSS distance (x-axis, log scale) across nine gene-relative annotation contexts (1–5 kb upstream, 5′UTR, promoter, first exon, 3′UTR, exon, CDS, intron–exon boundary, exon–intron boundary). Red lines represent SNPs residing inside ASM-DMRs and blue lines represent SNPs outside ASM-DMRs. Shaded bands indicate 95% confidence intervals. The number of SNP–gene pairs (N) is shown for each context in matching colors. **(E)** Correlation between ASM delta and eQTL NES (y-axis) as a function of relative SNP position to the DMR center (x-axis, DMR-centered with outside regions compressed), across the same nine gene-relative annotation contexts as in (**D**). Red lines show the running correlation with 95% confidence intervals (shaded). Gray shading marks the interior of the DMR (positions −10 to +10).

Having defined discrepancy between ASM and mQTL signals (**Figure 5A–B**), we next examined whether allele-specific methylation captures downstream regulatory effects on gene expression by comparing ASM direction and magnitude with GTEx eQTL signals. Concordance was defined based on the agreement between the direction of allele-specific methylation (DMR delta) and the direction of gene expression changes (GTEx prostate tissue eQTL normalized effect size, NES), such that concordant pairs reflect consistent regulatory effects, i.e., higher methylation associated with lower gene expression (**Figure 5A**). Using this framework, variants located within DMRs showed modest but significantly higher concordance with eQTL directionality compared to variants outside DMR (p = 0.028; **Figure 5C**), suggesting that SNPs residing within ASM-defined regions are more likely to reflect functional regulatory effects on gene expression. We next examined how concordance varies with SNP-to-TSS distance across genomic annotations (**Figure 5D**). Concordance showed a robust decay with increasing distance from transcription start sites, with the strongest agreement observed for variants proximal to TSS and progressively diminishing at distal regions. This pattern was consistent across most genomic features, indicating that proximity to canonical regulatory elements remains a strong determinant of expression-associated allele-specific epigenetic effects. To test whether the relative position of a SNP to DMR influences the allele-specific methylation to GTEx effect, we evaluated the correlation between allele-specific methylation (ASM delta) and GTEx (NES) as a function of SNP position relative to DMRs (**Figure 5E**). Interestingly, the correlation pattern was genomic-context specific: promoter- and 5UTR-associated DMRs showed consistently negative ASM–eQTL correlations across most relative positions, whereas 3UTR-associated DMRs displayed weaker positive correlation and in distant SNP-to-DMR positions.

### Hotspot ASM at the IRX4 locus nominates a candidate androgen-responsive regulatory element

The IRX4 locus has been previously reported as a prostate cancer GWAS risk locus and is also associated with gene expression variation (eQTL), indicating its potential regulatory role. After examining the ASM pattern in this region, we observed that an ASM cluster with larger methylation differences were concentrated within a relatively narrow genomic interval (chr5:1888662-1890105), while weaker signals were more dispersed (**Figure 6A**). To further characterize the regulatory context, we integrated AR ChIP–seq data (PRJNA752082) and found that this interval overlapped AR binding peaks that were higher in prostate adenocarcinoma than in normal prostate (**Figure 6B**). In Castration-Resistant Prostate Cancer (CRPC) tissue (PRJNA777879), AR binding at this site increased upon DHT treatment and was accompanied by elevated H3K27ac signal (**Figure 6C**), consistent with androgen-responsive regulatory activity. Notably, the local SNP structure lacks a clear LD block (**Figure 6D**), and ASM signals are observed across multiple nearby variants. Among these, rs6885084 shows a cancer-specific methylation difference, consistent with the cancer-associated changes in AR ChIP–seq profiles. This difference is further supported by read-level methylation patterns, which show a larger allelic methylation difference in cancer samples than in normal samples (**Figure 6E**), highlighting rs6885084 as a candidate variant for further functional investigation.

**Figure 6.**
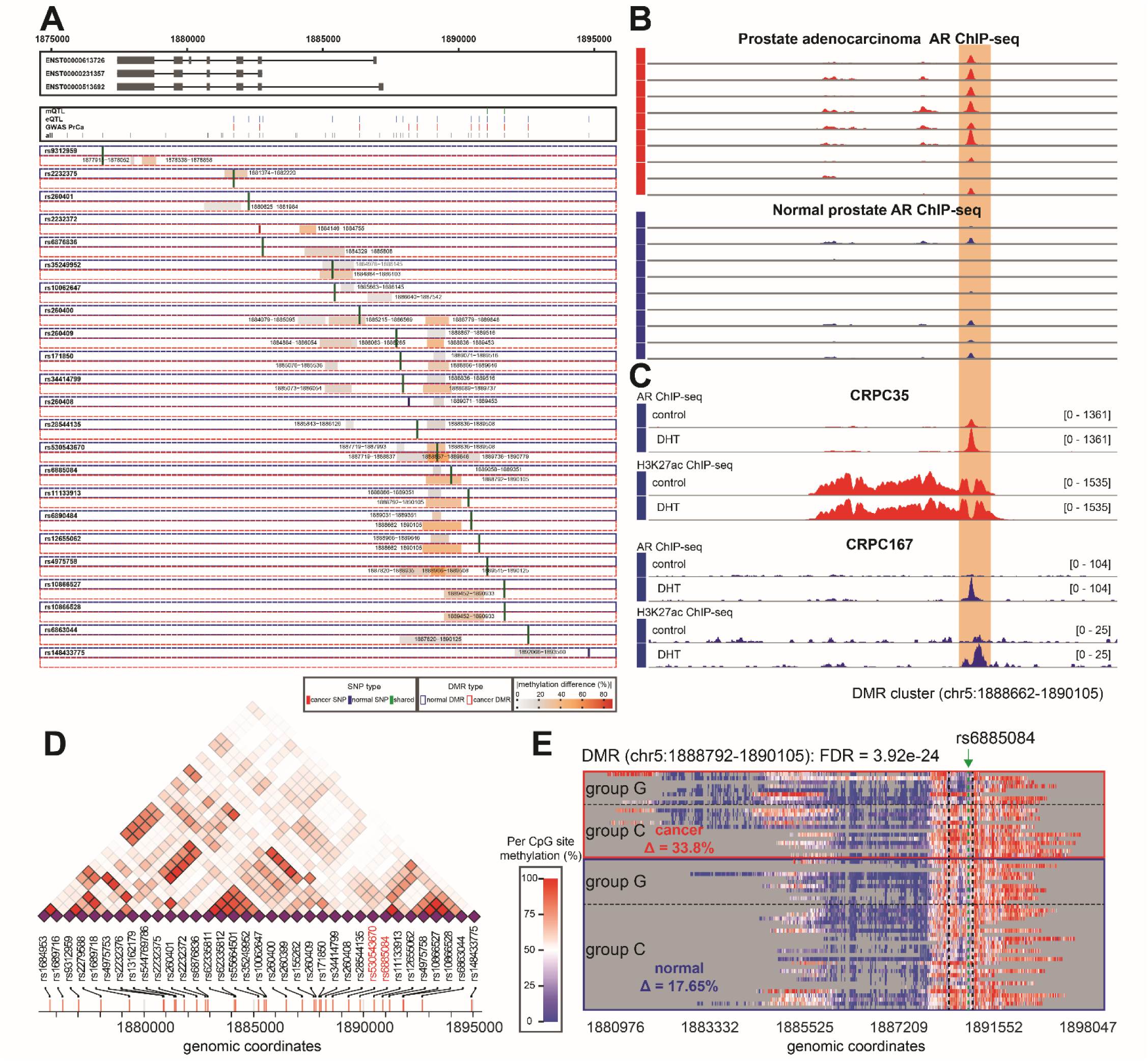
Case study of a prostate cancer GWAS locus on chromosome 5 with allele-specific methylation and androgen receptor binding. **(A)** Locus overview showing IRX4 gene models (top), mQTL, eQTL, and GWAS prostate cancer risk SNPs (colored ticks), and ASM-DMRs for each input SNP (rows). Each row corresponds to one SNP (rsID labeled on left); colored rectangles indicate ASM-DMR intervals, with fill color representing methylation difference (%) between alleles. Cancer SNPs are shown in red, normal SNPs in blue, and shared SNPs in green. Normal DMRs are outlined in blue and cancer DMRs in red. **(B)** AR ChIP-seq signal tracks from prostate adenocarcinoma (top, red) and normal prostate (bottom, blue) tissue samples at the same locus. The orange-shaded region (chr5:1,888,662–1,890,105) marks a DMR cluster overlapping a prominent AR binding peak that is substantially stronger in cancer than in normal tissue. **(C)** AR ChIP-seq and H3K27ac ChIP-seq tracks from two castration-resistant prostate cancer (CRPC) (CRPC35 and CRPC167) under vehicle control and DHT stimulation conditions. The orange-shaded DMR cluster region shows strong DHT-induced AR binding and H3K27ac enrichment, indicating androgen-responsive enhancer activation. **(D)** Linkage disequilibrium (LD) heatmap of all input SNPs across the locus, with r² values represented by color intensity (white = low LD, red = high LD). **(E)** Methylation heatmap for the lead ASM-DMR (chr5:1,888,792–1,890,105; FDR = 3.92×10⁻²O) stratified by allele at rs6885084 (G vs. C, green arrow). Each row represents the per-site methylation profile of one allele in one patient sample, with red indicating higher methylation and blue indicating lower methylation.

### The ASM-associated locus linked PSCA gene expression to rs4736369

PSCA gene is being evaluated as a therapeutic target in metastatic castration-resistant prostate cancer (mCRPC).^48^ We therefore examined whether allele-specific methylation (ASM) at this locus was associated with PSCA regulation. Across the tested variants, we identified a prominent ASM signal centered on rs4736369 in both normal and cancer samples (**Figure 7A**). The corresponding ∼500-bp DMR showed methylation differences of 50.7% and 53.4% in normal and cancer cohorts, respectively (**Figure 7B**). Although no significant eQTL signal was detected in the GTEx normal prostate cohort, multiple significant eQTL and isoform-QTL associations were observed in the MAYO prostate cohort (prostate cancer–adjacent normal tissue) (**Figure 7C**). Gene set enrichment analysis further indicated that PSCA expression was associated with multiple prostate-related pathways (**Figure 7D**). In sashimi plot, RNA-seq coverage profiles showed differences in junction usage between genotype groups, with higher coverage and junction reads observed in C/C compared to T/T samples (**Figure 7E**), consistent with alternative splicing at this locus. Leveraging long-read phasing information from nanopore sequencing, we performed haplotype-resolved analysis in heterozygous individuals. Allele directed haplotype-specific expression analysis, integrating nanopore DNA and RNA-seq data, showed that variants phased with the rs4736369 C allele are associated with higher RNA expression (**Figure 7F**). Consistently, ChIP–seq data (PRJNA540151) from heterozygous prostate tissue demonstrated higher H3K4me3 signal on the rs4736369 C allele than on the T allele (**Figure 7G**).

**Figure 7.**
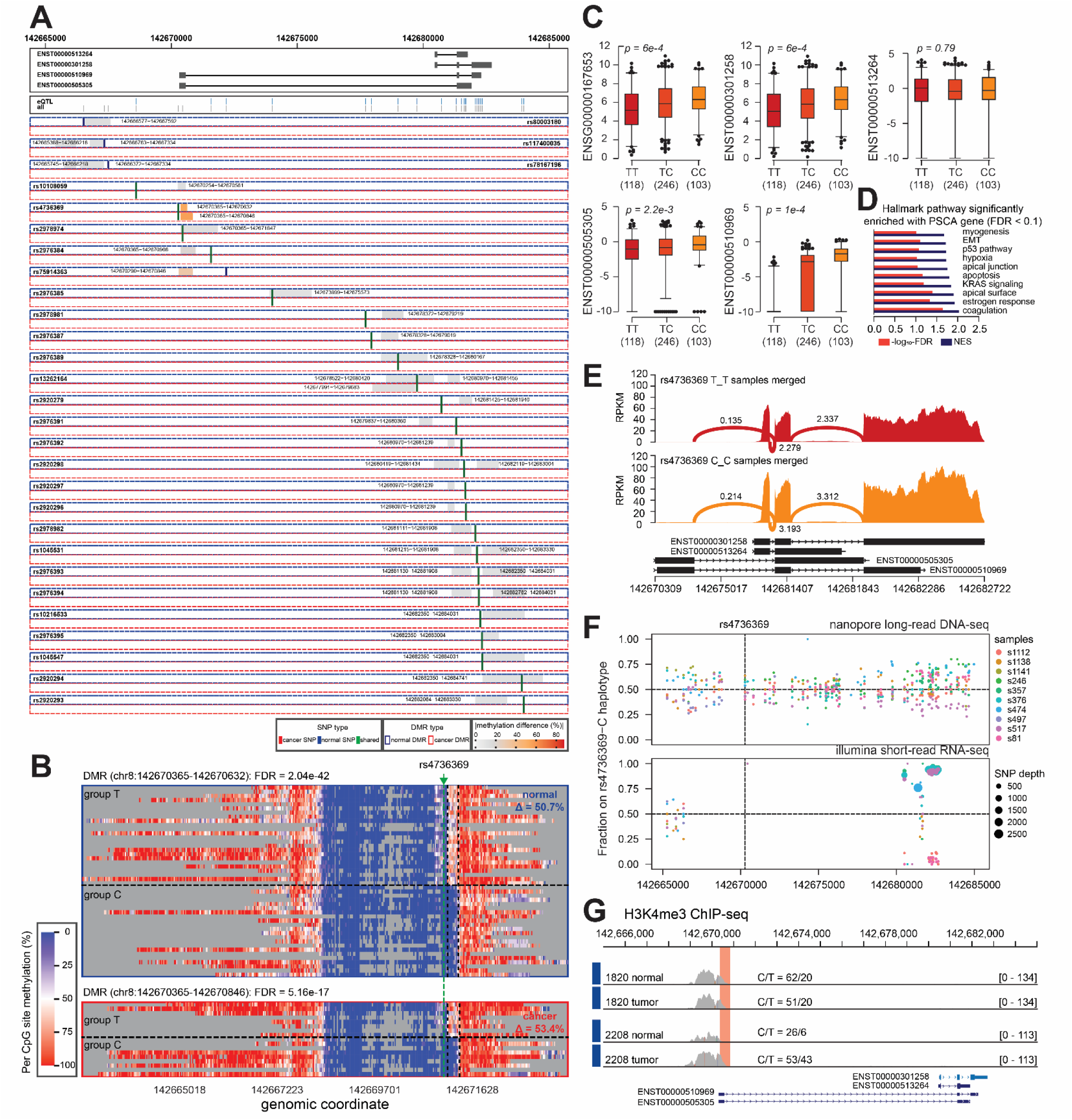
Allele-specific methylation at a prostate cancer GWAS locus on chromosome 8 links rs4736369 to transcriptional regulation. **(A)** Locus overview showing PSCA gene models (top), eQTL SNPs (blue ticks), and ASM-DMR intervals for each input SNP (rows). Each row corresponds to one SNP (rsID labeled on left); colored rectangles indicate ASM-DMR intervals with fill color representing methylation difference (%) between alleles. Cancer SNPs are shown in red, normal SNPs in blue, and shared SNPs in green. Normal DMRs are outlined in blue and cancer DMRs in red. The lead SNP rs4736369 (green arrow) is highlighted. **(B)** Per-CpG site methylation heatmaps for the ASM-DMRs at rs4736369 in normal and cancer tissue, stratified by allele group (T vs. C). Each row represents the per-site methylation profile of one allele in one patient sample; red indicates higher methylation and blue indicates lower methylation. **(C)** Boxplots of normalized gene expression for five transcripts at this locus (ENSG00000167653, ENST00000301258, ENST00000513264, ENST00000505305, ENST00000510969) stratified by rs4736369 genotype (TT, TC, CC; sample sizes in parentheses). **(D)** Hallmark pathway enrichment analysis for genes significantly associated with the PSCA eQTL (FDR < 0.1) at this locus. Bar length represents −log₁₀(FDR) (red) and normalized enrichment score (NES, blue). **(E)** Sashimi plots of RNA-seq read coverage and splice junction usage at the locus, generated from BAM files merged by rs4736369 genotype: homozygous T/T (top, red) and homozygous C/C (bottom, orange). Arc heights and labeled scores indicate splice junction read counts. **(F)** Haplotype-directed allele-specific RNA-seq coverage at rs4736369, where haplotype phasing was determined from nanopore long-read DNA sequencing. Each dot represents one Mayo Clinic sample, colored by sample ID, plotted as the allele fraction of the C allele at the rs4736369 position in DNA (top) and RNA-seq (bottom); dot size in the RNA-seq panel reflects SNP read depth. DNA allele fractions cluster near 0, 0.5, or 1.0, confirming germline genotype assignments. RNA-seq allele fractions show systematic deviation from 0.5 in heterozygous samples, indicating allele-specific expression consistent with the eQTL direction and the allele-specific methylation observed in (**B**). **(G)** H3K4me3 ChIP-seq tracks at the locus for matched normal and tumor tissue from two patients (1820 and 2208). The red-shaded region marks the ASM-DMR interval. Allele counts at rs4736369 (C/T) are indicated for each sample. H3K4me3 signal is present right near the DMR in both normal and tumor tissue, indicating that this region overlaps an active promoter, and the signal intensity varies between samples with different allele compositions, supporting allele-specific chromatin activity at this locus.

### Discussion

Nanopore sequencing uses deep neural networks with alignment-free decoding to directly infer nucleotide sequences and base modifications from raw ionic current signals, enabling robust measurement of regional DNA methylation. Leveraging these single-molecule, multimodal data, we first characterized genome-wide tumor–normal differential methylation regions and their genomic and functional features. Our whole-genome nanopore sequencing design enabled comprehensive profiling of cancer-associated methylation, revealing strong enrichment of DNA-binding and transcriptional regulatory functions. These findings support a model in which early-stage prostate cancer–associated hypermethylation preferentially targets CpG islands, whereas hypomethylation occurs more broadly across the genome.^49-52^ To further assess the relationship between DNA modification and transcriptional activity, we examined modification patterns across genes stratified by prostate tissue-specific expression levels derived from GTEx RNA-seq data. Both 5mCG and 5hmCG exhibited clear expression-dependent distributions, characterized by pronounced depletion around transcription start sites (TSS) and enrichment across gene bodies. These patterns suggest that DNA modifications may serve as surrogate markers of transcriptional activity. To this end, we explored the functional relevance of these observations by performing GSEA using gene-level aggregated 5hmCG signals. Consistent with previous reports ^53-55^, hallmark and chromosomal gene sets exhibited modestly higher 5hmCG levels in normal samples compared to tumor tissues. In contrast, enrichment analysis of positional gene sets (C1) revealed that mitochondrial genes showed significantly elevated 5hmCG levels in cancer relative to normal tissue, consistent with increased oxidative phosphorylation activity^56^ and mitochondrial metabolism^57^ observed in tumors. This distinct pattern may reflect reduced cellular heterogeneity and lower background variability in the mitochondrial genome compared to autosomal regions, thereby enabling more robust detection of modification differences even in small sample sizes. In addition to mtDNA copy number,^58,59^ mutation^60,61^ and 5mCG methylation,^62,63^ these observations raise the possibility that mitochondrial 5hmCG modification patterns could serve as potential biomarkers for tumor detection, particularly in liquid biopsy or circulating tumor DNA (ctDNA) settings.

Beyond per-site methylation measurements, long-read sequencing enables the analysis of fragment-level combinations of CpG sites, allowing quantification of regional methylation entropy. This fragment-level view captures coordinated methylation patterns and provides a measure of epigenetic heterogeneity that cannot be inferred from average methylation levels alone. Notably, conventional methylation entropy metrics are typically derived from unmodified cytosine and 5mCG states,^64,65^ whereas the entropy framework used in this study incorporates both 5mCG and 5hmCG signals, thereby capturing additional layers of epigenetic information. Contrary to the intuition that cancer epigenomes are more disordered, we observed a significant reduction in methylation entropy specifically within DMRs that gained methylation in cancer tissue, while entropy in hypomethylated or unchanged DMRs remained unaltered. The entropy reduction at cancer hypermethylated DMRs reflects the clonal fixation of a uniform 5mCG state across the tumor cell population — a process underpinned by two converging mechanisms. First, 5hmCG is broadly lost in cancer as a consequence of TET downregulation^66^ or dysfunction.^67,68^ Second, once 5hmCG is depleted, the remaining hemi-5mCG at replication forks is faithfully restored by DNMT1, which has a strong and well-characterized preference for hemi-methylated over hemi-hydroxymethylated substrates.^45^ The 5mCG state is therefore heritably propagated through cell division while 5hmCG is passively diluted, and the expanding tumor clone converges on a low-entropy, uniformly methylated pattern.^46,47^ Across different histone modifications, the highest entropy was co-located at enhancer markers, which are characterized by partial demethylation and enrichment of 5hmCG,^69,70^ generating a mixed modification landscape across reads. This confirms that entropy is not a linear marker of chromatin activity but specifically reports on modification heterogeneity, which is maximal at regulatory elements poised between methylated and demethylated states.

The structured methylation patterns revealed by expression-dependent analyses and entropy profiling suggest that epigenetic states are not purely stochastic but are shaped by underlying mechanisms, including local germline variation. To enable this analysis, we developed a simple allele-specific methylation (nanoASM) pipeline to systematically identify and quantify allele-specific methylation events from nanopore long-read sequencing data. A key assumption underlying this method is that functionally relevant germline variants exert local (cis) effects on the epigenetic landscape, preferentially influencing nearby CpG sites rather than distal regions. Although many mQTL studies^17,71-74^ have demonstrated predominantly cis-acting genetic effects of SNPs on nearby CpG sites, the distance thresholds used to define cis associations vary widely across studies, typically ranging from 250 kb to 1 Mb. This variability partly reflects inherent limitations of genotype-to–single-CpG association designs. We have previously discussed potential biases associated with this approach.^22^ In this study, we further observed that discrepancies between genetic and methylation signals were greater when SNPs were in proximity to the target CpG sites. This may be explained by probe-binding artifacts in array-based platforms, where SNPs within or near probe sequences can introduce allele-specific hybridization biases, thereby affecting methylation measurements. Similar effects have been reported previously.^20,21^

In our nanoASM framework, we propose an approach that aggregates allelic methylation signals across local 8–20 kb regions flanking germline variants, enabling robust detection of ASM and improving the identification of regulatory SNPs. Interestingly, although the germline SNPs were largely shared between normal and cancer tissues, the corresponding ASM signals were mostly not reproduced across conditions, indicating substantial differences in the epigenetic landscape between normal and cancer states. We found that part of this discrepancy could be attributed to loss of heterozygosity (LOH) in cancer, while a larger fraction likely reflects genuine differences in epigenetic regulation between normal and tumor tissues. Notably, cancer-derived ASM regions exhibited broader genomic span and larger methylation differences compared to normal tissues. This observation is consistent with previous next-generation sequencing (NGS) studies ^75-78^ and suggests that epigenetic dysregulation in cancer operates at a larger regional scale. Leveraging nanopore long-read sequencing, which preserves haplotype-resolved allele information, we further confirmed that SNPs disrupting CpG sites are associated with local hypomethylation of DMRs, whereas SNPs creating CpG sites are associated with regional hypermethylation.^79,80^

Another advantage of variant-level ASM analysis is that, compared to haplotype-based approaches, it reduces diffuse effects from imprinting regions and better isolates the local impact of individual alleles on nearby methylation patterns. ^81,82^. In addition, haplotype-based phasing can result in heterogeneous block structures across individuals, complicating cross-sample comparisons and may introduce inconsistency in statistical analyses.^83^ In contrast, variant-level analysis provides a consistent and well-defined unit of comparison across samples. Interestingly, in conventional mQTL analyses, we observed that SNPs located further away from CpG sites exhibit greater discrepancy between mQTL effect sizes and allele-specific methylation differences, compared to proximal SNP–CpG pairs. We hypothesize that this pattern is driven by two factors. First, long-range SNP–CpG associations are more likely to be influenced by LD, where the tested SNP may not be the causal variant, leading to allele switching and inconsistent effect directions.^84^ Second, mQTL estimates are often derived from single CpG measurements from array-based data, which are more susceptible to technical noise and site-specific variability, in contrast to allele-specific methylation that aggregates signals across reads and local CpG contexts.^85^

A central challenge in post-GWAS functional genomics is the identification of both the causal regulatory variant and the functional domain it controls, tasks that conventional association-based approaches struggle to resolve simultaneously due to LD confounding and the absence of direct functional readout ^86^. The two loci presented here illustrate two distinct but complementary utilities of variant-level ASM analysis in translational cancer epigenomics, highlighting it a unified framework to address both challenges. At the IRX4locus, the ASM-DMR anchored by rs6885084 delineates an androgen-responsive enhancer domain, evidenced by its precise colocalization with cancer-specific AR ChIP-seq occupancy and H3K27ac signal that is further potentiated by DHT stimulation in CRPC cell lines. This is consistent with the well-established enrichment of sequence-dependent ASM at active regulatory elements, including enhancers and transcription factor binding sites. ^82,87^ Importantly, the amplification of the allelic methylation difference in cancer tissue relative to normal tissue — a pattern documented more broadly across cancer types^77^ suggests that tumor-specific epigenetic remodeling does not create ASM de novo but rather potentiates a pre-existing constitutive cis-regulatory signal. At the PSCA locus, the analytical logic shifts from domain delineation to causal variant resolution. By examining allelic methylation differences, ASM analysis identifies rs4736369 as the likely causal cis-regulatory variant, a conclusion supported by the convergent evidence of allele-specific H3K4me3 occupancy, and allele-specific expression validated concordantly across nanopore long-read and Illumina short-read platforms. This exemplifies the use of ASM as a post-GWAS fine-mapping strategy, where the SNP anchoring the local allelic methylation difference is a cis-acting functional candidate — providing orthogonal, functionally grounded evidence that statistical fine-mapping methods based on LD alone cannot achieve. Taken together, the two loci establish that variant-level ASM analysis serves a dual purpose: it identifies the spatial boundaries of functional regulatory domains and resolves the identity of causal regulatory variants within them.

## Acknowledgements

This study has been supported by the National Institutes of Health [R01CA250018 and R01CA212097, to L.Wang. and R01CA263494 to L.Wu]. The funders had no role in study design, data collection and analysis, publication decision, or manuscript preparation.

## Author Contributions Statement

Conception and design: Y. Tian, J. Wong, S. McDonnell, N. Larson, L. Wu, B. Manley, L. Wang

Development of methodology: Y. Tian, J. Wong, S. McDonnell, N. Larson, L. Wu, B. Manley, L. Wang

Sample curation and collection: B. Manley.

Acquisition of data: Y. Tian, J. Wong,

Analysis and interpretation of data: Y. Tian, J. Wong, H. Zhong, L. Wu, S. McDonnell, N. Larson

Writing, review, and revision of the manuscript: Y. Tian, J. Wong, S. McDonnell, N. Larson, H. Zhong, L. Wu, B. Manley, L. Wang

Study supervision: Y. Tian, N. Larson, L. Wu, B. Manley, L. Wang

## Conflicts of Interest Statement

L.Wu. provided consulting service to Pupil Bio Inc. and reviewed manuscripts for Gastroenterology Report, not related to this study, and received honorarium. No potential conflicts of interest were disclosed by other authors.

---

**Figure S1.**
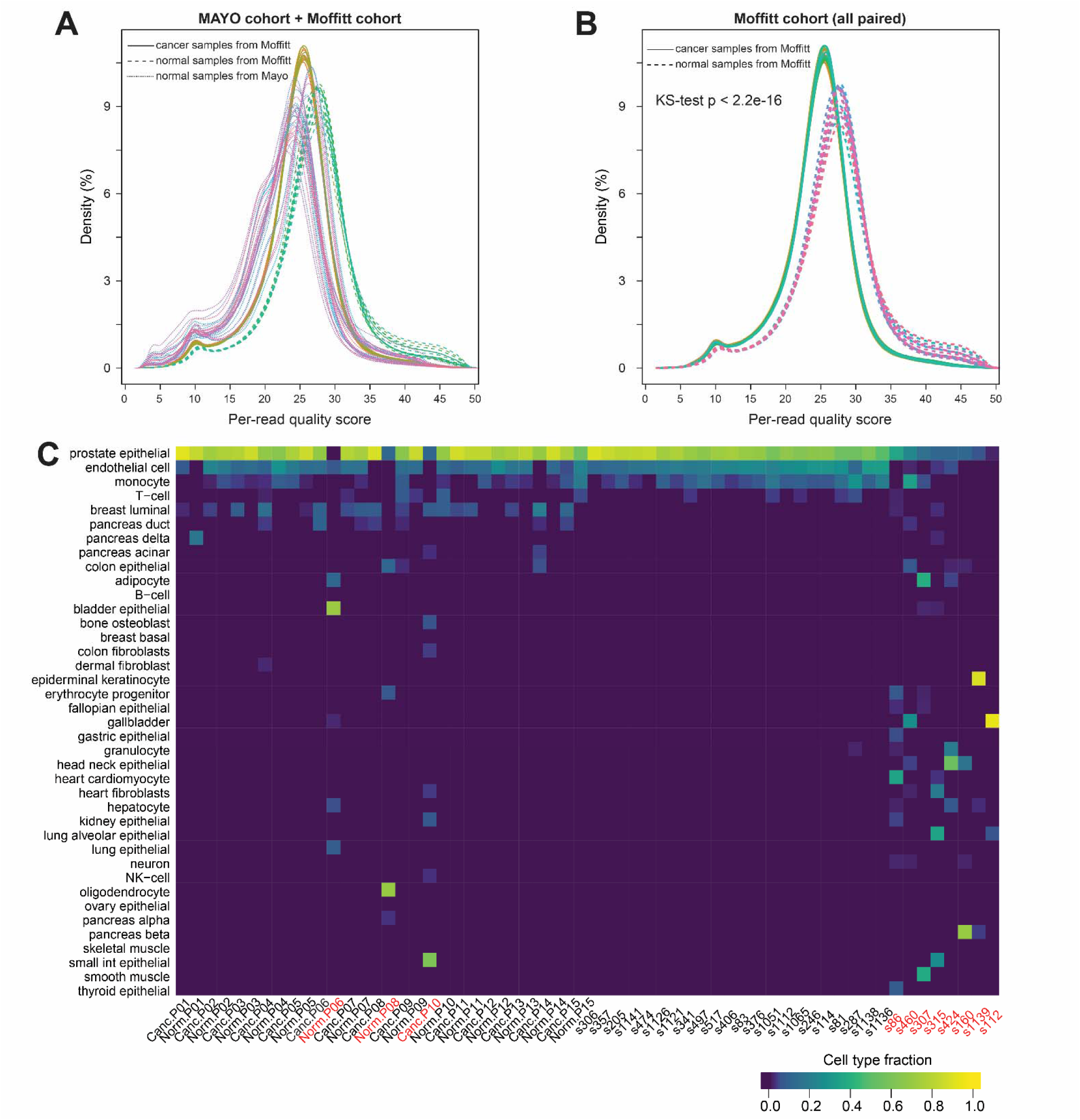
Nanopore sequencing quality control and cell-type deconvolution. **(A)** Per-read quality score distributions for all samples across the Mayo Clinic and Moffitt cohorts. Each curve represents one sample; solid lines indicate Moffitt cancer samples, dashed lines indicate Moffitt normal samples, and dotted lines indicate Mayo Clinic normal samples. The majority of reads in all samples achieve quality scores ≥Q20, consistent with expected R10.4.1 PromethION performance. **(B)** Per-read quality score distributions restricted to paired tumor and normal samples from the Moffitt cohort. Solid lines represent cancer samples and dashed lines represent matched normal samples. Cancer samples exhibit a systematic leftward shift relative to paired normal samples (Kolmogorov–Smirnov test, p < 2.2×10⁻¹O), indicating lower overall basecalling quality in tumor-derived DNA. **(C)** Cell-type deconvolution heatmap showing the estimated fraction of 39 reference cell types across all samples, inferred from nanopore-derived CpG methylation profiles using the ATLAS reference panel. Prostate epithelial cells (top row) dominate the majority of samples. Samples with insufficient prostate epithelial fractions or low purity ratios, highlighted in red on the x-axis, were excluded from downstream analyses (Moffitt: Cancer.P10, Normal.P06, Normal.P08; Mayo Clinic: s86, s460, s307, s160, s315, s424, s1139, s112).

**Figure S2.**
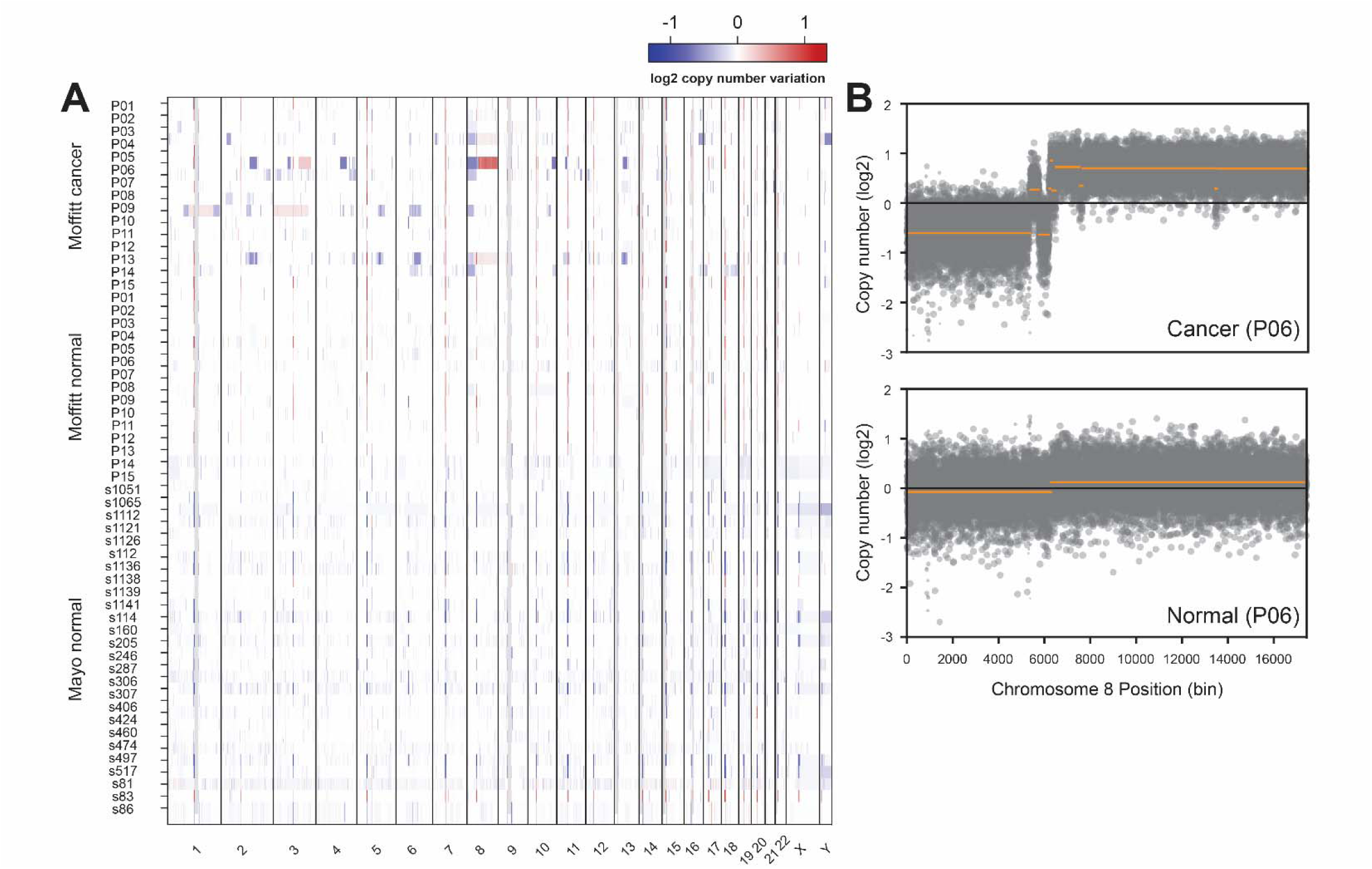
Genome-wide copy number variation analysis of prostate tissues. **(A)** Sample-wise heatmap of log₂ copy number ratios across all autosomes and sex chromosomes for Moffitt cancer (top), Moffitt normal (middle), and Mayo Clinic normal (bottom) samples. Each row represents one sample and each column represents a chromosomal region. Red indicates copy number gain and blue indicates copy number loss relative to the pooled normal reference. Moffitt cancer samples exhibit substantially greater copy number variation compared with normal tissues from either cohort. **(B)** Log_2_ copy number ratio scatter plots across chromosome 8 bins for the matched cancer (top) and normal (bottom) tissue from patient P06. Orange horizontal lines indicate CNVkit-derived segmentation calls. The cancer sample displays the characteristic prostate cancer pattern of 8p arm loss (bins ∼0–5500, segment below 0) and 8q arm gain/amplification (bins ∼6000–17000, segment above 0), which is absent in the paired normal tissue.

**Figure S3.**
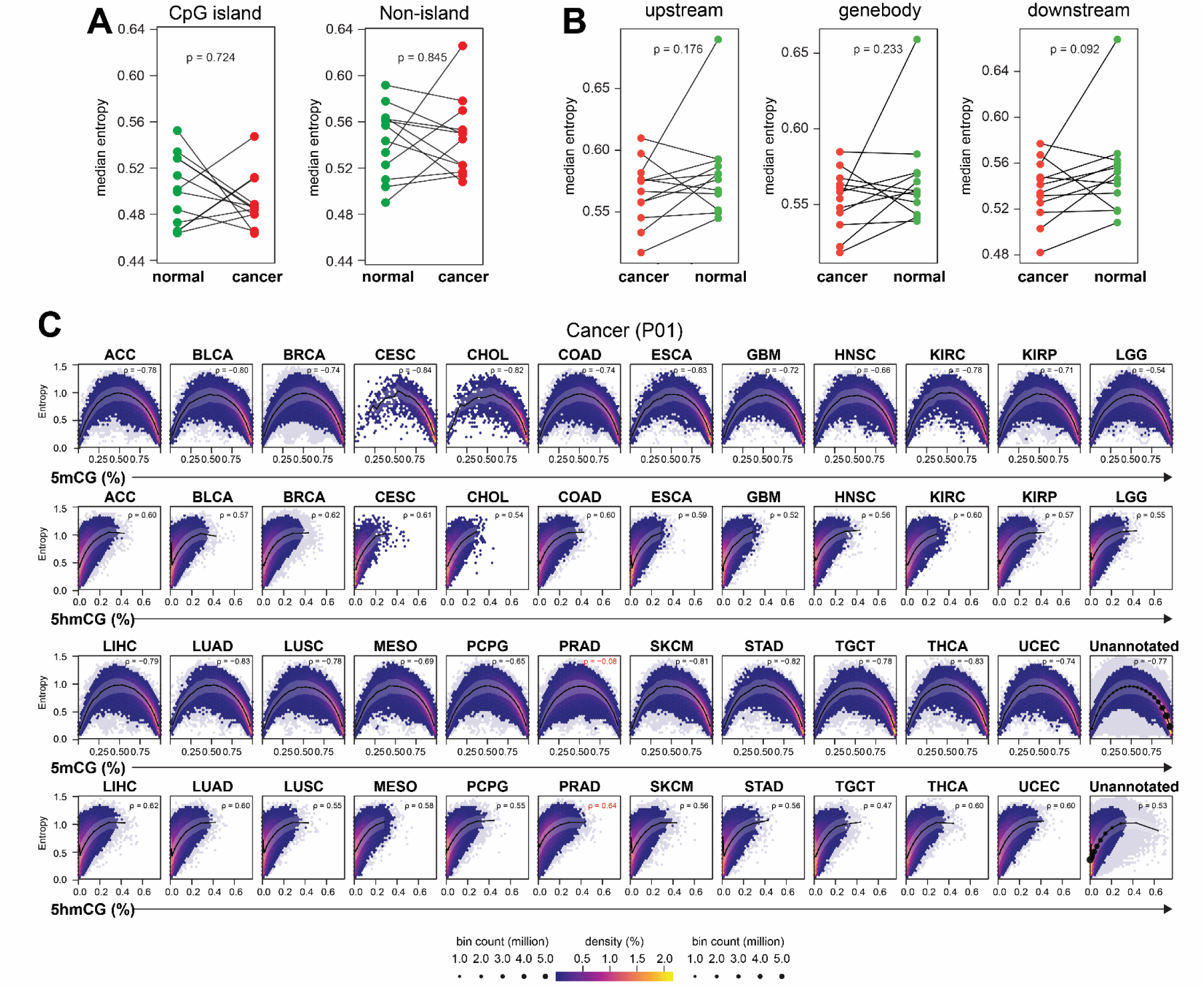
Methylation entropy analysis across genomic contexts and pan-cancer TCGA validation. **(A)** Paired dot plots comparing regional median methylation entropy between matched normal (green) and cancer (red) Moffitt samples, stratified by CpG island context: CpG island regions (left, p = 0.724) and non-island regions (right, p = 0.845). Lines connect paired samples from the same patient. **(B)** Paired dot plots comparing regional median methylation entropy between matched cancer (red) and normal (green) Moffitt samples across three gene-relative positional contexts: upstream (left, p = 0.176), gene body (middle, p = 0.233), and downstream (right, p = 0.092). Lines connect paired samples from the same patient. **(C)** Hexbin density scatter plots of methylation entropy versus 5mCG percentage (odd rows) and 5hmCG percentage (even rows) for the Moffitt cancer sample from patient P01, evaluated across 24 TCGA pan-cancer chromatin state annotations (ACC, BLCA, BRCA, CESC, CHOL, COAD, ESCA, GBM, HNSC, KIRC, KIRP, LGG, LIHC, LUAD, LUSC, MESO, PCPG, PRAD, SKCM, STAD, TGCT, THCA, UCEC, and Unannotated). Color scale indicates hexbin density (bin count in millions, left) and density percentage (right). Spearman correlation coefficients (ρ) are shown for each panel; values highlighted in red indicate a significant departure from the pan-cancer background pattern.

**Figure S4.**
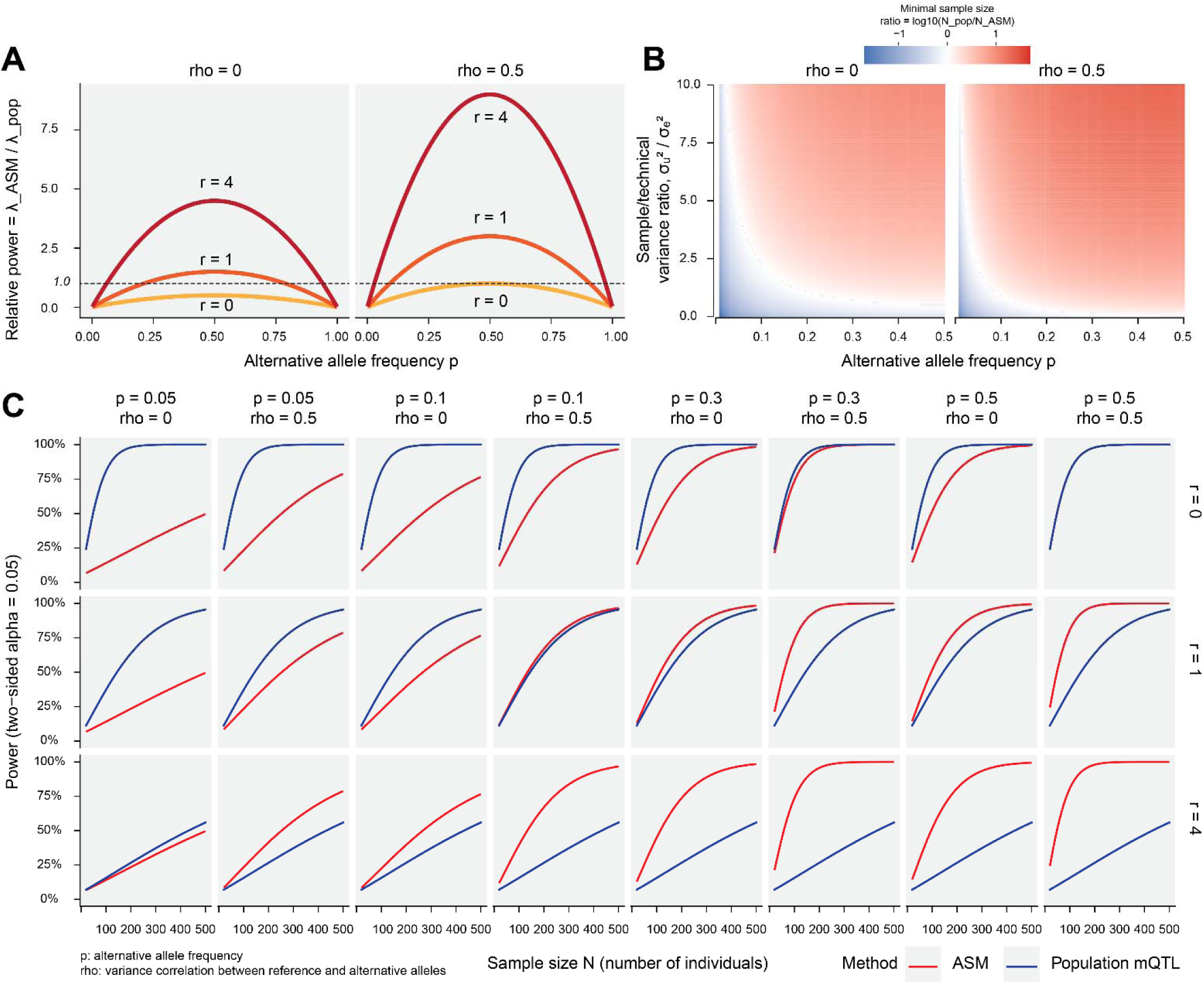
Theoretical power comparison between allele-specific methylation (ASM) and population-level mQTL analysis. **(A)** Relative statistical power of ASM versus population mQTL analysis (λ_ASM / λ_pop) as a function of alternative allele frequency (p), shown for two levels of within-individual allelic variance correlation (rho = 0, left; rho = 0.5, right) and three levels of sample/technical variance ratio (r = 0, orange; r = 1, red; r = 4, dark red). The dashed horizontal line at 1.0 marks the threshold of equivalent power between the two methods. ASM analysis achieves greater relative power than population mQTL across a broad range of allele frequencies, with the advantage increasing with higher variance ratio (r) and intermediate allele frequencies. When ρ = 0.5, the power advantage of ASM is substantially amplified, particularly for common variants (p = 0.3–0.5). **(B)** Heatmap of the minimal sample size ratio (log₁₀(N_pop / N_ASM)) required to achieve equivalent power between population mQTL and ASM analysis, as a function of alternative allele frequency (p, x-axis) and sample/technical variance ratio (σ_u² / σ_e², y-axis), for rho = 0 (left) and rho = 0.5 (right). Red indicates that population mQTL requires a larger sample size than ASM (i.e., ASM is more efficient); blue indicates the reverse. Across most parameter combinations, ASM requires substantially fewer samples than population mQTL to achieve equivalent power, with the greatest efficiency gains at intermediate allele frequencies and high variance ratios. **(C)** Power curves (two-sided α = 0.05) as a function of sample size N (number of individuals, x-axis) for ASM (red) and population mQTL (blue), across a grid of alternative allele frequencies (p = 0.05, 0.1, 0.3, 0.5; columns), allelic variance correlations (ρ = 0 and ρ = 0.5; column pairs), and variance ratios (r = 0, top row; r = 1, middle row; r = 4, bottom row). ASM consistently achieves higher power than population mQTL at equivalent sample sizes across most parameter combinations, with the largest gains at higher variance ratios (r = 4) and intermediate allele frequencies. The power advantage of ASM is further amplified when rho = 0.5, reflecting the benefit of the shared within-individual genetic background in the allele-resolved design.

