## Supplementary material for "nanoASM: Long-Read Allele-Specific DNA Methylation Profiling Enables Functional Annotation of Regulatory Noncoding Variants in Human Prostate Tissues": NoteS1

**Methods and materials**

**Statistical Model and Theoretical Power Analysis for Allele-Specific Methylation–Based cis-mQTL Detection**

To compare the statistical efficiency of allele-specific methylation (ASM)–based cis-mQTL testing with conventional population-level mQTL analysis, we formulated a simple random-effects model for DNA methylation measurements at SNP-associated regions. This model captures two primary sources of variability: (i) sample-level heterogeneity (e.g., cell-type composition, donor background, and batch effects), and (ii) measurement noise arising from methylation estimation.

**Random-effects model for population-level methylation**

Let $Y_{i}$denote the bulk methylation level for individual $i$, and $G_{i}$the genotype dosage. We consider the model:

$$Y_{i}=\mu+\beta G_{i}+U_{i}+\varepsilon_{i},$$

where $G_{i}\in\{0,1,2\}$ is genotype dosage (ALT allele count), $\beta$ is the population-level mQTL effect, $U_{i}\sim N(0,\sigma_{U}^{2})$ represents sample-level variability, and $\varepsilon_{i}$captures measurement noise. To provide a consistent comparison with allele-specific measurements, we define bulk methylation as the average of allele-specific quantities:

$$Y_{i}=\frac{Y_{i,\mathrm{ref}}+Y_{i,\mathrm{alt}}}{2}.$$

Under this definition, the technical variance of $Y_{i}$is reduced relative to allele-level measurements, yielding:

$$\mathrm{Var}(Y_{i})=\sigma_{U}^{2}+\frac{\sigma_{e}^{2}}{2}.$$

Thus, population-level mQTL detection must resolve genetic effects in the presence of both sample-level heterogeneity and measurement noise.

For convenience, we define the variance ratio

$$r=\frac{\sigma_{U}^{2}}{\sigma_{e}^{2}},$$

which quantifies the relative contribution of sample-level variability to technical noise.

**ASM model for heterozygous individuals**

For heterozygous individuals, allele-specific methylation can be estimated separately:

$$Y_{i,\mathrm{ref}}=\mu+U_{i}+\varepsilon_{i,\mathrm{ref}},Y_{i,\mathrm{alt}}=\mu+\delta+U_{i}+\varepsilon_{i,\mathrm{alt}},$$

where $\delta$represents the true allelic methylation difference. Defining the within-sample contrast:

$$D_{i}=Y_{i,\mathrm{ref}}-Y_{i,\mathrm{alt}},$$

we obtain:

$$D_{i}=-\delta+(\varepsilon_{i,\mathrm{ref}}-\varepsilon_{i,\mathrm{alt}}),$$

and therefore:

$$\mathrm{Var}(D_{i})=2\sigma_{e}^{2}(1-\rho),$$

where $\rho=\mathrm{Corr}(\varepsilon_{i,\mathrm{ref}},\varepsilon_{i,\mathrm{alt}})$, which captures the extent to which measurement noise is shared between the two allele-specific methylation estimates. Because both alleles are derived from the same biological sample and processed through identical experimental workflows, including DNA extraction, library preparation, sequencing, and base modification calling, their technical errors are not expected to be independent. In the limiting case where technical errors are perfectly correlated ($\rho\to1$), the variance of the allelic contrast approaches zero, whereas when errors are independent ($\rho=0$), the variance reduces to the baseline value of $2\sigma_{e}^{2}$.Importantly, the sample-level term $U_{i}$cancels exactly, making ASM a paired measurement that is robust to inter-individual variability.

**Noncentrality parameters and statistical power**

Under large-sample approximations, statistical power is governed by noncentrality parameters (NCPs):

Population-level test:

$$\lambda_{\mathrm{pop}}=\frac{N\beta^{2}}{\sigma_{e}^{2}(r+1/2)}$$

ASM-based test:

$$\lambda_{\mathrm{ASM}}=\frac{4N\beta^{2}p(1-p)}{\sigma_{e}^{2}(1-\rho)}$$

where $r=\sigma_{U}^{2}/\sigma_{e}^{2}$, and $p$is the allele frequency.

**Relative statistical efficiency: Power ratio** $\boldsymbol{R(p)}$**comparing ASM and mQTL**

The ratio of statistical efficiency is:

$$R(p)=\frac{\lambda_{\mathrm{ASM}}}{\lambda_{\mathrm{pop}}}=4p(1-p)\cdot\frac{r+\frac{1}{2}}{1-\rho}$$

Interpretation

This formulation highlights several key features:

- ASM gains power by eliminating sample-level variability through within-individual contrasts.
- The efficiency advantage increases with allele frequency, peaking near $p=0.5$.
- Larger sample-level variability (higher $r$) further favors ASM.
- Positive correlation between allele-specific technical errors ($\rho>0$) reduces the variance of the ASM contrast, further improving efficiency.

**Threshold allele frequency and sample size scaling**

ASM is expected to outperform population-level testing when:

$$4p(1-p)\cdot\frac{r+\frac{1}{2}}{1-\rho}>1$$

This condition is most readily satisfied for common variants and datasets with substantial sample-level heterogeneity. To achieve a fixed power level, the required sample sizes scale as:

$$N_{\mathrm{pop}}\propto(r+\frac{1}{2}), N_{\mathrm{ASM}}\propto\frac{1-\rho}{4p(1-p)}$$

Thus, the relative efficiency directly translates into differences in required cohort size.

**Simulation-based illustrations (Figures S1–S3)**

- Figure S1A shows $R(p)$across allele frequencies and variance ratios, illustrating the dependence of efficiency on alternative allele frequency, signal-to-noise ratio and variance of allele-specific methylation measurements.
- Figure S1B presents heatmaps of minimal sample size ratios required to achieve 80% power.
- Figure S1C shows power as a function of sample size under different parameter settings.

**Practical implications**

1. ASM removes sample-level heterogeneity through paired allelic contrasts.
2. Efficiency gains are strongest for common variants with sufficient heterozygous carriers.
3. In realistic datasets, where sample-level variability is substantial, ASM can provide meaningful gains in statistical efficiency.
4. The magnitude of improvement depends on allele frequency, variance structure, and technical error correlation.

**Conclusion**

Under a general random-effects framework, ASM-based cis-mQTL testing provides improved statistical efficiency by eliminating shared sample-level variation and reducing effective noise in paired measurements. These advantages are most pronounced for common variants and heterogeneous datasets, supporting the use of ASM as a primary strategy for detecting cis-regulatory methylation effects.
